# Evolution of mechanical chromatin insulators from selfish genetic elements

**DOI:** 10.64898/2026.08.08.743475

**Authors:** Faisal Alkhaldi, Monika Priyadarshini, Sonia El Mouridi, Hajar Al-Zarah, Satoshi Habuchi, Christian Frøkjær-Jensen

## Abstract

Genomes require physical boundaries to separate active and repressed chromatin, a function traditionally attributed to insulator proteins. Here, we show that genomic insulation can also emerge from physical properties encoded directly within the DNA polymer. In *C. elegans*, transcription-coupled mutational bias remodels Helitron transposon minisatellites to match the DNA helical repeat. The resulting 10-bp periodicity (PATCs) encodes intrinsic curvature, creating topologically responsive DNA elements that favor local deformation under supercoiling, disrupt canonical nucleosome organization, and protect germline genes from progressive and heritable silencing. Rather than acting as short protein-binding motifs, Helitron-derived PATCs form extended barrier elements that limit the stabilization of repressive chromatin. Comparative analyses suggest that related sequence-encoded mechanical signatures recur across Metazoa, including at *Drosophila* insulators, human CTCF sites, and active human LINE-1 retrotransposons. Thus, sequence-encoded mechanics provide an evolutionarily accessible substrate for chromatin insulation—a physical layer of genome organization that can be co-opted by host genomes to protect gene expression, and potentially retained by selfish elements for their own persistence.

## Introduction

A genome is a long, confined DNA polymer whose sequence encodes not only genetic information, but also the energetic cost of mechanical deformation [1, 2]. Sequence-encoded mechanics can therefore shape the 3D folding and organization of chromosomes. For example, phased A-tracts encode intrinsic curvature at replication origins [3], poly(dA:dT) tracts encode local stiffness that resists nucleosome wrapping [4], and (TG/CA)*^n^* repeats encode torsional compliance that buffers supercoiling stress [5]. These mechanical codes are typically distributed and quantitative [6]: their effects are weak at the scale of a few base pairs and strengthen only when integrated over tens to hundreds of base pairs or repeated copies. Unlike protein-binding motifs that arise from local sequence changes [7], mechanical function is a distributed property, raising the question of how such extended patterns evolve to bias chromosome topology.

Nowhere is this mechanical requirement more acute than in chromatin insulation and the partitioning of the genome into regulatory domains. Unlike enhancer-blocking, which regulates distant interactions, the barrier activity of insulators provides a physical blockade against the linear spreading of histone modifications. These barriers allow regulatory activity and chromatin state to be controlled locally. In many animals, insulating boundaries are positioned by the architectural protein CTCF [8]. However, *C. elegans* lacks a canonical CTCF-like insulator system [9], or any other known insulators, yet its autosomes are partitioned into alternating chromatin domains enriched for H3K27me3 and H3K36me3, that impose strong expression rules in the germline [10]. Germline transgenes in repressive domains are frequently silenced, yet this repression is gradual across generations, reversible, and temperature-sensitive, implying that domain insulation relies on a dynamic physical barrier rather than a static regulatory state [11, 12]. One clue to how insulation can be achieved without canonical boundary proteins comes from Periodic A_N_/T_N_ Clusters (PATCs). PATCs are long, AT-rich, non-coding elements with a ∼10-bp periodicity that have been identified by anomalous gel migration, distinctive topological conformations, and bioinformatic analysis of the *C. elegans* genome, implying a sequence-encoded structure that reflects unusual DNA mechanics [13, 14]. PATCs can license germline expression of transgenes within repressive chromatin [15, 16] and were proposed to protect endogenous genes from piRNA-mediated silencing [17]. Together, this motivated the hypothesis that PATCs act as a genomic watermark—a signal that allows surveillance pathways to distinguish endogenous “self” genes from foreign DNA, including transposable elements, and protect them from potent germline silencing mechanisms [15, 17]. Despite their prevalence in the *C. elegans* genome and their rapid gain on nematode evolutionary timescales [15], at least two basic questions remain unresolved: what evolutionary process drives the emergence of this specific 10-bp periodicity, and how does this feature enforce a chromatin barrier?

Here, we identify an evolutionary origin and functional logic for these elements. We trace PATCs to the co-option of ancient Helitron DNA transposons [18], identifying a transcription-coupled mutational process that remodels internal minisatellites to match the DNA helical repeat. The resulting 10-bp periodicity encodes intrinsic curvature and promotes a compact, nucleosome-disruptive DNA structure that is consistent with the preferential localization of torsional stress. Functionally, this evolved periodicity protects germline expression in repressive chromatin and limits the persistence of transgenerational silencing. We therefore propose that Helitron-derived PATCs evolved as sequence-encoded chromatin barriers that help maintain gene-expression homeostasis in the *C. elegans* germline. More broadly, comparative analyses suggest that related mechanical signatures recur at insulator-associated and transposon-derived sequences across animals, including human CTCF boundaries and active LINE-1 retrotransposons.

## Results

### Helitron minisatellites are the source of PATCs

The evolutionary source of PATCs has remained elusive, despite their rapid turnover and positive correlation with repressive chromatin environments [15]. To identify their origin, we quantified the genome-wide overlap between PATC density and annotated genomic features [19, 20]. This analysis revealed that the vast majority of the PATC signal (∼80%) co-localizes with Helitron transposable elements (Fig. 1a, Supplementary Fig. S1a), specifically the non-autonomous families HelitronY1A and HelitronY1 (Supplementary Fig. S2a). The remaining PATC signal was not annotated as Helitron-derived and included smaller contributions from other repeat classes as well as non-transposable-element regions. This association is unexpected, raising the question of how a transposon—typically a target of silencing—came to encode a genomic ‘self’ signal.

**Figure 1:**
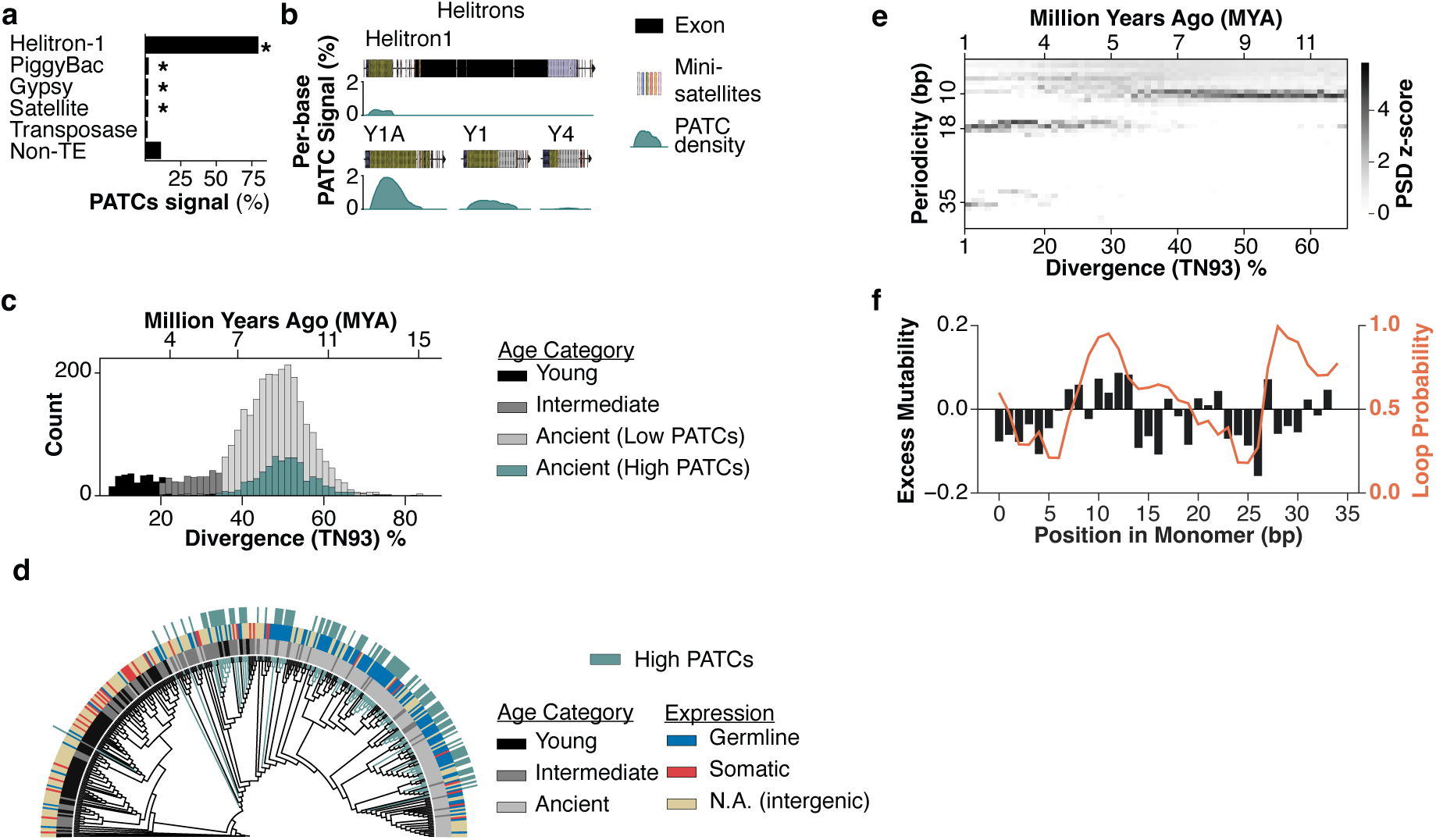
Helitron minisatellites are the evolutionary source of PATCs. **a,** Overlap of genomic PATC signal with annotated repeat classes (One-sided, chromosome-preserving interval-relocation permutation tests were performed using 10,000 permutations: Helitron-1, P = 1.00 × 10−4, n = 8,635; PiggyBac, P = 0.00730, n = 9,314; Gypsy, P = 1.00 × 10−4, n = 881; Satellite, P = 1.00 × 10−4, n = 8,101; Transposase, P = 1.00, n = 22,281. An asterisk denotes P < 0.05, and n denotes annotated repeat elements.). **b,** Genomic PATC density mapped onto Helitron consensus sequences, identifying internal minisatellites as the signal source. **c,** Age distribution of Helitron-derived minisatellites estimated by nucleotide divergence (TN93). Colors indicate classification into Young, Intermediate, or Ancient (High/Low PATC) categories. **d,** Phylogenetic tree of Helitron-derived minisatellites stratified by age category, host gene expression and PATC density. The tree has been randomly sub-sampled for visual clarity. **e,** Evolutionary spectrogram of Helitron minisatellite periodic structure versus sequence divergence (TN93). Heatmap intensity indicates PSD z-score. **f,** Per-position excess mutability (black bars, left axis) and predicted ssDNA loop formation probability (red line, right axis) across the 35-bp monomer.

We next investigated whether this signal is intrinsic to the transposon or acquired after insertion, by comparing the PATC density of individual genomic Helitron insertions against their ancestral consensus sequences (Supplementary Fig. S1b, Supplementary Fig. S2b). While genomic copies frequently exhibit high PATC scores, the consensus sequences for all major families are devoid of the signal. Mapping the cumulative genomic PATC signal onto the Helitron consensus reveals that this difference is spatially confined: the PATC signal maps specifically to an internal 35-bp minisatellite repeat within Helitron1 and its non-autonomous derivatives, particularly HelitronY1A and HelitronY1 (Fig. 1b, Supplementary Fig. S1b).

To determine the timing and context of this acquisition, we reconstructed the evolutionary history of the PATC-precursor minisatellites by using a profile Hidden Markov Model (HMM) and sequence divergence as a molecular clock. This analysis dates the majority of PATC-bearing Helitron insertions to a major expansion approximately 7–11 million years ago (Fig. 1c). Stratifying these ancient insertions by host gene expression reveals that high PATC density evolved predominantly in Helitrons within germline-active genes (comprising ubiquitous and germline-specific transcripts), whereas those in soma-specific genes remained PATC-poor (Supplementary Fig. S2c).

Finally, to test whether High PATC states arose once or recurrently, we constructed a sequence-similarity tree using the HMM-derived ancestral 35-bp minisatellite consensus as the root (Fig. 1d). High PATC elements did not form a single monophyletic branch, but instead appeared across multiple branches of the tree. Consistent with recurrent acquisition, fixed-root parsimony mapping from a PATC-poor ancestral state inferred 363 Low-to-High transitions and 79 High-to-Low transitions. Thus, PATCs are not ancestral Helitron features, but derived DNA structure that repeatedly accumulate after genomic insertion, especially in germline-associated contexts. This recurrent and context-biased acquisition motivated us to ask what mutational process could repeatedly remodel the ancestral 35-bp minisatellite into a 10-bp periodic sequence.

### Transcription-coupled bias drives evolution

Identifying the ancestral sequence offers a unique opportunity to reconstruct the evolutionary emergence of a distributed sequence feature. We quantified the periodicity of Helitron-derived minisatellites across their evolutionary lifespan using Power Spectral Density (PSD) analysis (Fig. 1e). The resulting spectrogram reveals a striking shift in dominant periodicity: young elements retain the ancestral ∼35-bp repeat, which decays into an intermediate 16–18-bp signal before converging on a robust 10-bp periodicity in ancient elements. This trajectory indicates that the 10-bp feature is an emergent property of minisatellite evolution rather than an ancestral feature. Furthermore, the systematic convergence of independent loci on this specific outcome implies that a widespread mutational bias drives this evolution.

Given that mutation rates are often governed by local sequence context [21], we reasoned that the systematic convergence could arise from position-specific mutational biases within the fundamental 35-bp monomer. We therefore developed a “monomer-folding” alignment strategy that maps individual repeat units onto a common 35-bp coordinate system (Supplementary Fig. S3a). This approach allows us to aggregate thousands of independent mutational events, revealing the precise distribution of mutation rates across the ancestral monomer. To distinguish functional remodeling from neutral drift, we calculated an “excess mutability” score that isolates mutations specific to the high PATC lineage (Fig. 1f), revealing a distinct position-specific signature. Given the strict restriction to germline-active genes (Supplementary Fig. S2c), we hypothesized a transcription-coupled mechanism. Transcription is known to transiently expose single-stranded DNA (ssDNA) in repetitive sequences, enabling formation of secondary structures that drive repeat instability and mutational bias [22]. Consistent with this, excess mutability correlates positively with the propensity for predicted ssDNA secondary-structure formation across the monomer (Spearman’s ρ = 0.406, P = 0.0155, n = 35 monomer positions; Fig. 1f). This suggests that repeat-intrinsic folding creates mutational hotspots during germline transcription, driving heritable remodeling toward the observed periodicity.

To test if this signature is sufficient to generate the PATC-defining 10-bp periodicity, we performed forward evolutionary simulations starting from the inferred ancestral minisatellite sequence. Applying the inferred high PATC mutational rates reproducibly produced sequences with the 10-bp periodic signal (Supplementary Fig. S2d; Supplementary Fig. S4), whereas simulations driven by the low PATC profile, or initiated from a shuffled ancestral sequence, did not. Given the low probability of achieving high PATC scores by chance [14], these results confirm that the inferred transcription-coupled mutational bias is sufficient to drive the emergence of PATCs from the ancestral Helitron.

### Ongoing remodeling in wild populations

Our model predicts that PATC accumulation should be an ongoing process driven by germline transcription. To test this, we examined Helitron-derived minisatellites across 17 wild *C. elegans* isolates [23–25]. While the minisatellite loci themselves are highly stable (Supplementary Fig. S5a)—predominantly representing ancient, fixed insertions—their internal PATC density varies. Comparing orthologous elements reveals that PATC density increases significantly in germline-active genes relative to somatic or intergenic contexts (Fig. 2a, Supplementary Fig. S5b, P < 0.001). Thus, natural variation in wild isolates supports a model in which PATC evolution remains active within fixed Helitron-derived minisatellites and is biased by germline transcription.

**Figure 2:**
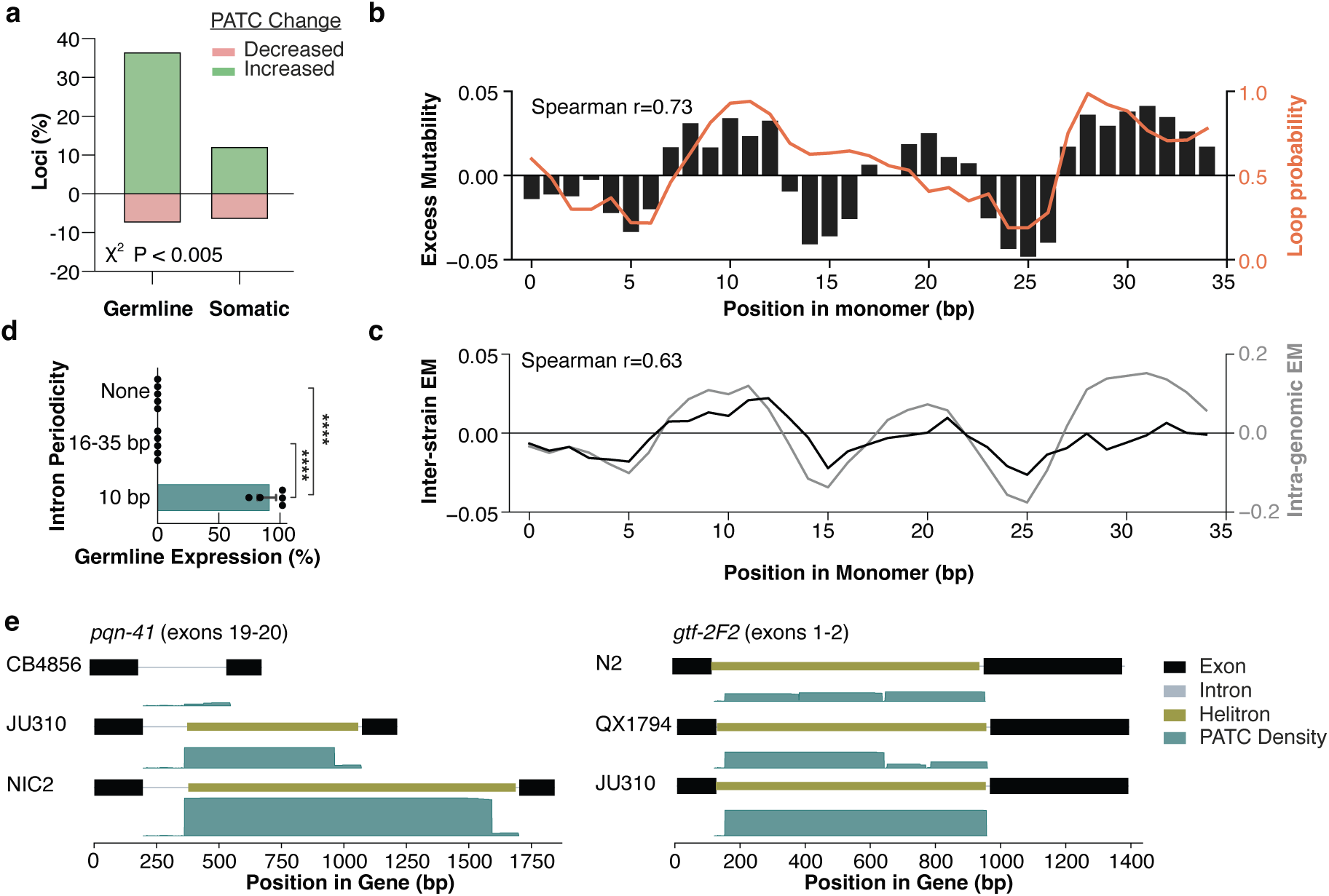
Ongoing germline-specific evolution of PATCs in wild populations. **a**, Change in PATC density for conserved loci relative to N2 reference, categorized by host gene expression (Pearson χ^2^ test of independence: χ^2^(2) = 1,013.03, P = 1.06 × 10*^−^*^220^, n = 23,439 locus–isolate observations (germline, n = 19,144; somatic, n = 4,295). **b**, Excess mutability profile derived from inter-strain polymorphisms and predicted ssDNA loop formation probability (red line, right axis) across the 35-bp monomer (Two-sided Spearman rank correlation: ρ = 0.727, P = 7.40 × 10*^−^*^7^, n = 35 monomer positions.). **c**, Comparison of intra-genomic (black) and inter-strain (gray) excess mutability profiles (Two-sided Spearman rank correlation: ρ = 0.631, P = 4.76 × 10*^−^*^5^, n = 35 paired monomer positions.). **d**, Germline GFP expression in transgene silencing assay using reporters with expression- and length-matched controls (“none”, pCFJ2411), ancestral minisatellites (16-35 bp periodicity, pFMK38), or evolved sequences (10- bp periodicity, pCFJ2375) inserted into the *oxTi173* (V) locus. Dots represent independent lines (N = 5 per group); bars indicate mean (Two-sided unpaired Student t-tests (control versus 10-bp and 16–35-bp versus 10-bp): t(8) = −15.81, P = 2.56 × 10*^−^*^7^ for each comparison). **e**, Polymorphic loci *pqn-41* (left) and *gtf-2F2* (right) showing lineage-specific variation in minisatellites insertions and PATC signals.

To ask whether the same mutational process is still detectable in contemporary populations, we computed a monomer-resolution mutability profile using only inter-strain polymorphisms among wild isolates (Fig. 2b). Unlike the intra-genomic analysis above, which captures accumulated divergence from the ancestral consensus over millions of years, this comparison measures standing variation among orthologous minisatellites. The inter-strain profile closely matches the ancient intra-genomic pattern (Fig. 2c; Spearman’s *ρ* ≈ 0.63, P = 4.8 × 10*^−^*^5^), indicating that present-day polymorphisms preferentially occur at similar monomer positions remodeled during ancient PATC evolution. Moreover, inter-strain excess mutability is strongly correlated with predicted ssDNA secondary-structure formation probability across the 35-bp monomer (Spearman’s *ρ* = 0.73, P = 7.4 × 10*^−^*^7^, Fig. 2b). This association is stronger than that observed for the ancient intra-genomic divergence profile (Δ*ρ* = 0.33, bootstrap 95% CI: 0.60 to 0.09), consistent with contemporary polymorphisms providing a more direct readout of the active mutational process. Thus, the position-specific bias inferred from ancient Helitron remodeling is recapitulated in extant natural variation and is linked to ssDNA exposure within the repeat.

This ongoing evolution is visible at individual germline-expressed loci. For example, a Helitron-derived minisatellite insertion in *pqn-41* is absent in CB4856 but present in other isolates, where it displays lineage-specific variation in size and PATC density (Fig. 2e). Even when the insertion itself is fixed, as in *gtf-2F2*, it shows substantial divergence in PATC density. These examples confirm that Helitron-derived minisatellite remodeling is not a static historical event, but an ongoing, lineage-specific process in natural populations.

### Evolved periodicity confers insulation

In *C. elegans*, transgenes inserted into repressive chromatin environments are robustly silenced in the germline, whereas transgenes with PATC-rich introns can maintain germline expression from these otherwise silencing-prone loci [15]. This phenotype is analogous to classical barrier-insulator assays, in which a sequence is tested by its ability to protect an integrated reporter from position-effect repression [26]. We therefore asked whether the evolved 10-bp periodicity of Helitron-derived PATCs is sufficient to confer this protective activity, or whether insulation is instead a property of the ancestral minisatellite.

To test this, we engineered single-copy GFP reporters with introns containing either evolved 10-bp elements, their ancestral (∼35-16 bp) counterparts, or length-matched controls, and inserted them into a repressive genomic locus [15]. While all transgenes were expressed in the soma, only those containing the evolved 10-bp periodicity maintained germline expression (Fig. 2d). In contrast, transgenes carrying ancestral elements were silenced as effectively as length-matched controls. Thus, the protective activity is not an intrinsic property of the ancestral minisatellite, nor a generic enhancer feature of germline introns [16], but a gain-of-function adaptation conferred by the emergent 10-bp periodicity.

### Evolved periodicity encodes topologically responsive DNA curvature

The 10-bp periodicity of PATCs aligns A/T-tracts with the helical pitch, a motif that generates strong intrinsic curvature [2, 27]. Under torsional stress, DNA bending, twist, and writhe are physically coupled, allowing sequence-dependent curvature to bias where supercoiling is accommodated [28]. This coupling is particularly relevant to chromatin insulation, because transcription-generated supercoiling, topoisomerase activity, and DNA tension have been implicated in the formation and regulation of chromatin boundaries [29–31]. Since PATCs represent an extreme form of intrinsic curvature [13, 32], we hypothesized that the evolved 10-bp periodicity converts Helitron-derived minisatellites into topologically responsive DNA elements (TRDE), allowing sequence periodicity to bias local DNA structure under supercoiling (Fig. 3a).

**Figure 3:**
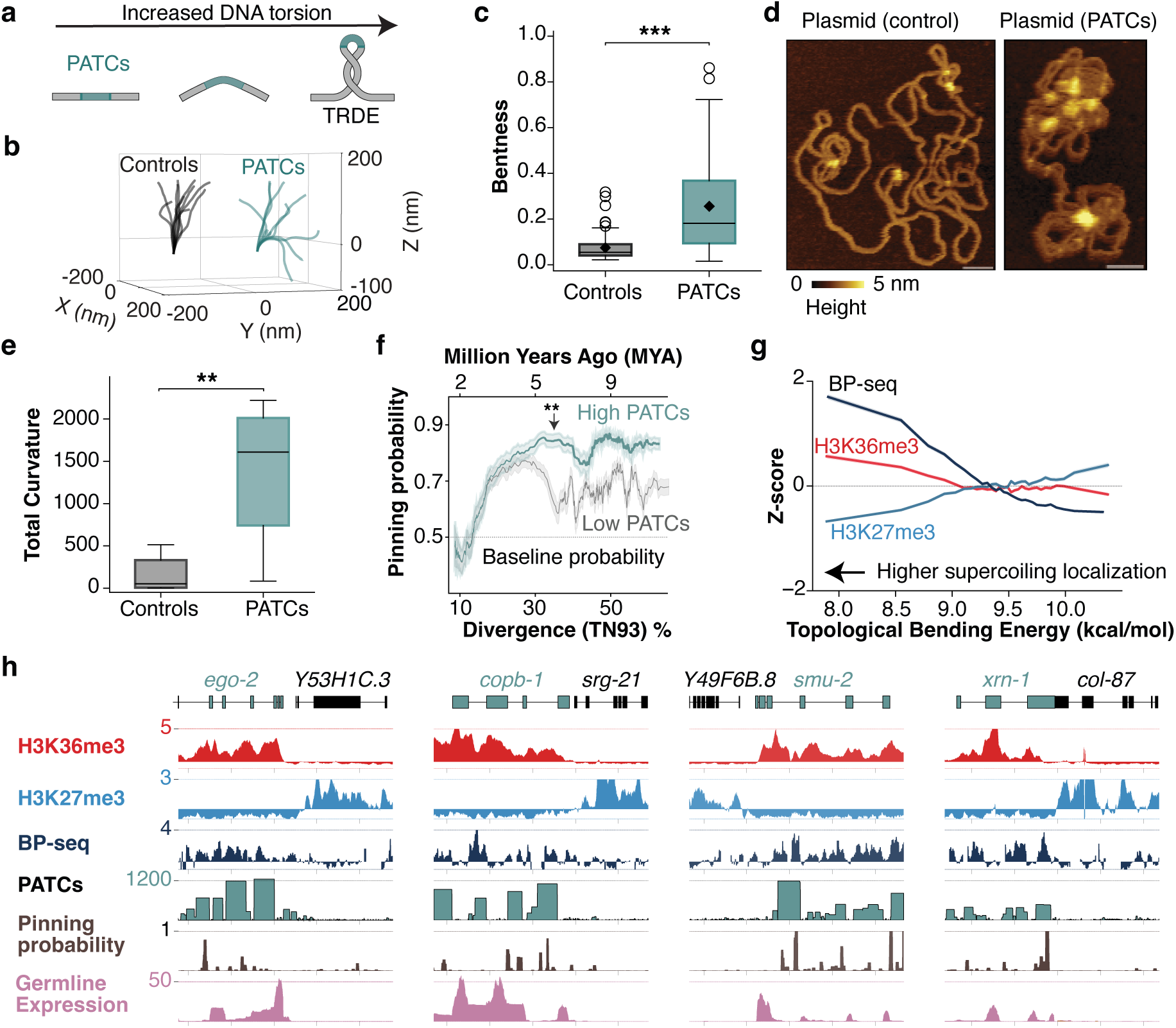
PATCs encode intrinsic curvature to pin supercoiling. **a**, Schematic of TRDE at the intrinsically bendable PATC-rich DNA sequence. **b**, Relaxed 3D conformations from sequence-dependent modeling for length-matched control (black) and PATC-rich (teal) introns (N = 10 per group, random genome sampling). **c**, Quantification of bending. Bentness (1 - (end-to-end distance / contour length)); (Two-sided Mann–Whitney U test: U = 1,591, P = 8.20 × 10*^−^*^17^, n = 100 sequences per group.). **d**, Liquid-phase AFM images of supercoiled control (pCFJ2411) and PATC-rich (pCFJ2375) plasmids. Scale bars: 100 nm. **e**, Total plasmid backbone curvature from dry AFM images (Following 1.5×IQR outlier exclusion, an exact two-sided Mann–Whitney U test gave U = 10 and P = 0.00216 (control, n = 12; PATC, n = 8; initial n = 12 and 10, respectively). **P < 0.01.). **f**, Pinning probability versus minisatellite divergence (TN93: bottom axis, MYA: top axis). Traces show high PATC versus low PATC lineages. Line thickness represents PATC density. Asterisks (**) indicate first statistically significant divergence (Two-sided bootstrap tests compared High and Low PATC groups across 18 TN93 bins (3,000 resamples; Benjamini–Hochberg correction). The earliest significant bin was centered at 29.54% TN93: Δmean = 0.130, P = 6.67 × 10*^−^*^4^, q = 0.00120 (Low PATC, n = 167; High PATC, n = 14). **q < 0.01.). **g**, Previously published genomic assays (BP-seq[36], H3K36me3 and H3K27me3[41]) versus predicted topological bending energy. **h**, Genome browser tracks (10-kb window) showing spatial alignment of previously published embryonic H3K36me3 and H3K27me3 [41], BP-seq [36], PATC density, pinning probability, and germline RNA-seq [41].

To quantify this structural bias, we modeled the conformational ensemble of Helitron-derived introns using a sequence-dependent statistical mechanics model [33]. In relaxed states, PATC-rich introns collapse into compact, highly curved structures, whereas length-matched controls remain extended (Fig. 3b). This difference is significant (Fig. 3c, P < 0.001), confirming that the evolved periodicity imposes a strong bending bias on the double helix.

We next asked if this bias alters DNA conformation under torsional stress. Closed circular DNA serves as a direct assay because its fixed linking number forces excess twist to be converted into writhe [34, 35]. We therefore visualized PATC-rich plasmids versus matched controls using atomic force microscopy (AFM). In liquid phase, PATC-containing plasmids adopted compact, highly writhed conformations with frequent self-contacts, contrasting with the open topologies of matched controls (Fig. 3d). Quantification of backbone curvature confirmed that PATC-rich plasmids exhibit significantly higher total curvature (Fig. 3e, P < 0.01). Together, these results show that the sequence-encoded bending bias alters DNA topology under constraint, favoring compact, writhed conformations.

To formalize this mechanical bias, we developed a quantitative Topological Bending Energy (TBE) model. The model asks a simple physical question: if torsional stress forces DNA to absorb sharp local bending, which sequences can accommodate that deformation at the lowest energetic cost? For each genomic window, we used a sequence-dependent statistical mechanics framework to estimate the relaxed shape and stiffness of the DNA, then calculated the energy required to impose a sharply bent geometry relevant to supercoiled and writhed DNA states [28, 33]. This approach builds on single-molecule measurements showing that DNA sequence can localize supercoils, and that intrinsic curvature is a major determinant of where plectonemic structures (interwound, rope-like supercoils) are localized on a naked DNA molecule [28].

The resulting TBE score represents an energy: lower values indicate that a sequence is already curved, or mechanically predisposed, in a way that makes local deformation easier. We then converted these energy values into a relative topological pinning probability using a Boltzmann weighting scheme. Pinning probability is therefore not a separate physical property, but a probabilistic interpretation of the same energy landscape: it estimates how likely a sequence is to capture supercoiling-induced deformation relative to other sequences in the genome. In this sense, TBE measures the cost of deformation, whereas pinning probability measures the expected localization bias produced by that cost difference. This metric accurately recapitulates experimentally measured supercoil-localization profiles from independent studies (Spearman’s ρ = 0.76; Supplementary Fig. S6) [28]. Applying the model to reconstructed Helitron minisatellite lineages revealed that high PATC evolution is accompanied by increased topological pinning probability (Fig. 3f), linking the emergent periodicity to local DNA deformation under torsional stress.

We next asked whether this sequence-intrinsic bias persists in the nucleus, where factors like nucleosomes and topoisomerases might override intrinsic mechanics. We analyzed several previously published genome-wide datasets; because these datasets come from different stages and conditions, we do not treat them as if they all came from the same developmental stage. Instead, we ask whether PATC-rich or low-TBE regions show enrichment within each dataset, using matched intron controls where appropriate. To connect TBE predictions to in vivo chromatin, we analyzed previously published genome-wide datasets, beginning with maps of negative supercoiling (BP-seq) generated by psoralen crosslinking [36]. This lets us test whether the same patterns recur across assays and stages, rather than being restricted to a single developmental context. Remarkably, predicted bending energy anti-correlates strongly with *in vivo* supercoiling (Spearman’s ρ = −0.65; Fig. 3g, Supplementary Fig. S6), indicating that intrinsic mechanics remain associated with topological structure in cells. This bias tracks with chromatin state: low-energy sequences are enriched in active domains (H3K36me3), while high-energy sequences coincide with repressive domains (H3K27me3) (Fig. 3h). Together, these results support a model in which the evolved 10-bp periodicity converts Helitron-derived minisatellites into mechanically biased DNA elements whose structural effects are amplified under torsional stress and remain detectable in chromatin.

### PATCs act as topological insulators

The properties of PATCs suggest a mechanism of insulation that differs from classical, sequence-specific insulators like CTCF. Rather than functioning as short, focal protein-binding elements positioned at domain borders, PATCs are long, intronic, and often distributed across kilobases within gene bodies [14]. To examine how these elements are positioned relative to chromatin domains, we mapped PATC density relative to the boundaries between active (H3K36me3) and repressive (H3K27me3) chromatin domains [10]. On autosomal arms, where repressive chromatin is pervasive, PATCs and low-TBE sequences are enriched not at focal boundary points, but across a broad transition zone extending several kb into the permissive chromatin region (Fig. 4a). This pattern is absent in the generally permissive autosomal centers (Supplementary Fig. S7), consistent with their distinct chromatin landscape [37] and no requirement for PATCs in transgenes to license germline expression [15].

**Figure 4:**
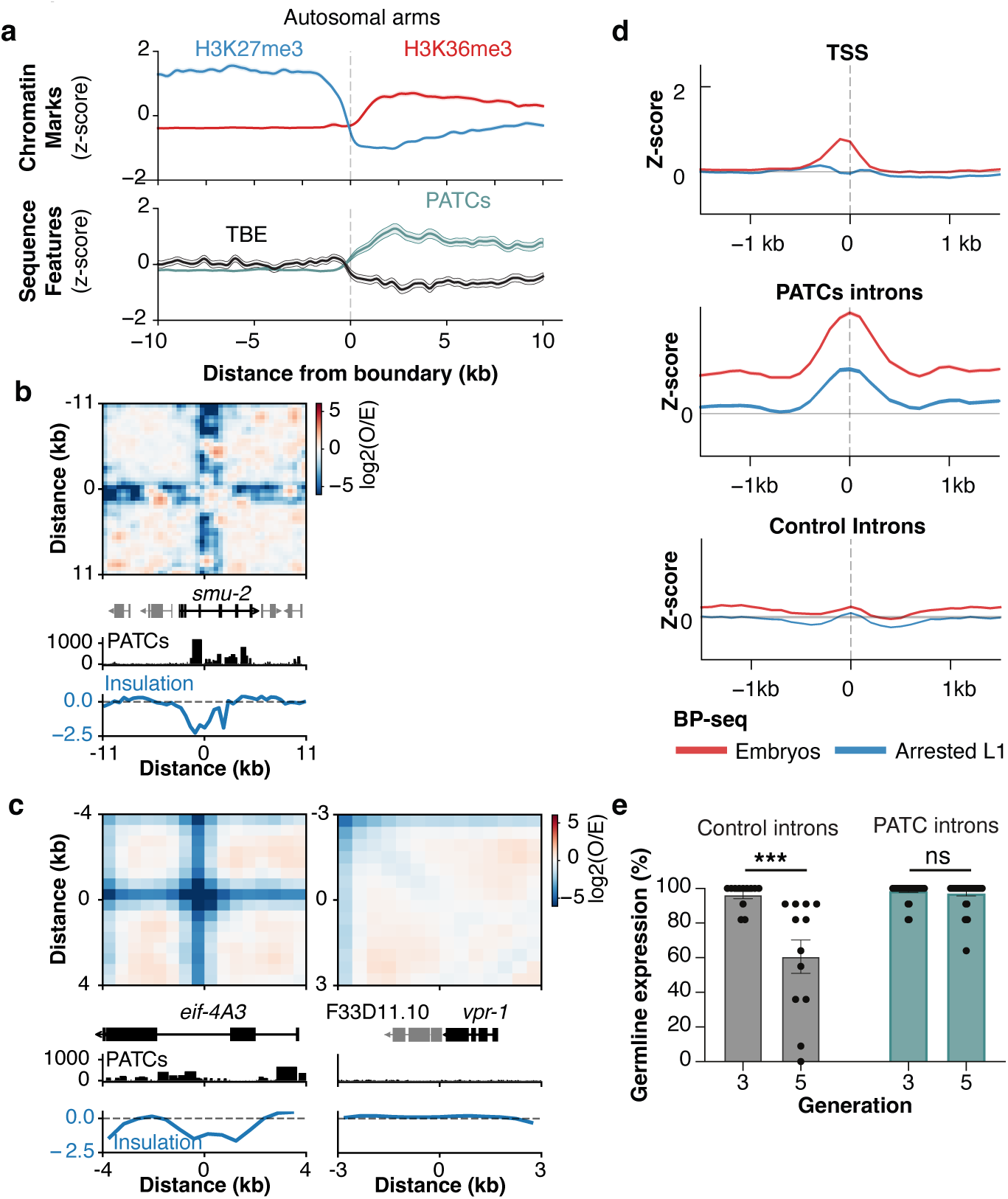
PATCs localize to chromatin boundaries and function as insulators. **a**, Previously published chromatin marks (ChIP-seq) [41] and sequence features aligned to autosomal-arm chromatin boundaries (x-axis: distance from boundary in kb). (**B-C**) Previously published Hi-C [38, 39] log_2_(observed/expected) view of *smu-2* locus (B) and the paralogs *eif-4A3* (left) and F33D11.10 (right, C). **d**, Previously published BP-seq [36] Z-scores for embryos (red) and arrested L1 (blue) aligned to transcription start sites (TSS), PATC-rich introns, and control introns. **e**, Transgenerational assay showing percentage of germline expression for control and PATC-containing (P*eft-3* ::*gfp*) reporters inserted into the *oxTi173* V repressive site. Control introns (P = 0.001, n = 12), PATC introns (P = 0.4802, n = 36), Mann-Whitney test.

We next asked whether PATCs coincide with local chromatin insulation at individual loci. In previously published high-resolution Hi-C maps [38, 39], the PATC-rich *smu-2* locus shows a local insulation feature aligned with PATCs (Fig. 4b). A paralog comparison further supports this association: the distal autosomal gene *eif-4A3*, which contains intronic Helitron/PATC insertions, shows a distinct local insulation boundary that is absent at the central autosomal PATC-poor paralog F33D11.10 (Fig. 4c). Aggregate Hi-C pileups show the same boundary-associated insulation pattern genome-wide, including after cohesin depletion (Supplementary Fig. S7).

This boundary localization is accompanied by localized topological stress: the previously published BP-seq [36] signal is significantly enriched over introns with high PATC density relative to controls (Fig. 4d). Notably, this enrichment persists in arrested L1 larvae even as the TSS-centered signals fade, confirming that PATC-associated supercoiling is a stable, intrinsic feature independent of ongoing transcription.

In the *C. elegans* germline, repression is not simply a static property of genomic position. Foreign or poorly optimized transgenes are often initially expressed [12], but then progressively acquire heritable silencing across generations as small-RNA and chromatin-based pathways reinforce the repressed state [40]. This provides a dynamic test of barrier function: if PATCs act as structural barriers to repressive chromatin, they should not only permit expression at a silencing-prone locus, but prevent an initially active transgene from being converted into a heritably silent state. We therefore monitored transgenerational expression of matched single-copy reporters inserted into a repressive domain. Control reporters lacking PATCs were initially active, but underwent progressive germline silencing within two generations (Fig. 4e). In contrast, PATC-containing reporters maintained robust germline expression. Thus, PATCs protect against the transgenerational establishment of germline silencing, supporting a model in which sequence-encoded DNA structure limits the propagation and stabilization of repressive chromatin across generations. This places PATCs at the interface between sequence-encoded DNA structure and epigenetic inheritance: they do not simply mark active chromatin, but prevent repressive states from being established.

### PATCs disrupt local nucleosome organization

The ability of PATCs to resist repressive chromatin suggests that they form a physical barrier to the nucleosome array. To determine the structural nature of this barrier, we analyzed the chromatin landscape of PATC-rich introns using previously published MNase-seq, ATAC-seq [41], and H3 CUT&RUN [42] datasets. Unlike Transcription Start Sites (TSS) which are accessible, or control introns which are nucleosome-occupied, PATC-rich sequences display a unique signature: a profound depletion of nucleosomes and H3 (Fig. 5d), accompanied by enrichment of “supra-nucleosomal” fragments in both MNase and ATAC assays (Fig. 5a, Supplementary Fig. S8a,b). These large fragments are resistant to digestion; in MNase titrations, they become progressively enriched as digestion intensity increases (Fig. 5b, Supplementary Fig. S8c), a result we validated using long-read Nanopore ATAC-seq to rule out mappability artifacts (Fig. 5c). Together, these assays identify PATC-rich introns as nucleosome-depleted but digestion-resistant regions, consistent with a compact non-nucleosomal structure that limits enzyme access. This footprint does not distinguish intrinsic DNA folding, torsionally constrained DNA, associated protein factors, or a combination of these mechanisms, but indicates that PATCs locally disrupt canonical nucleosome organization.

**Figure 5:**
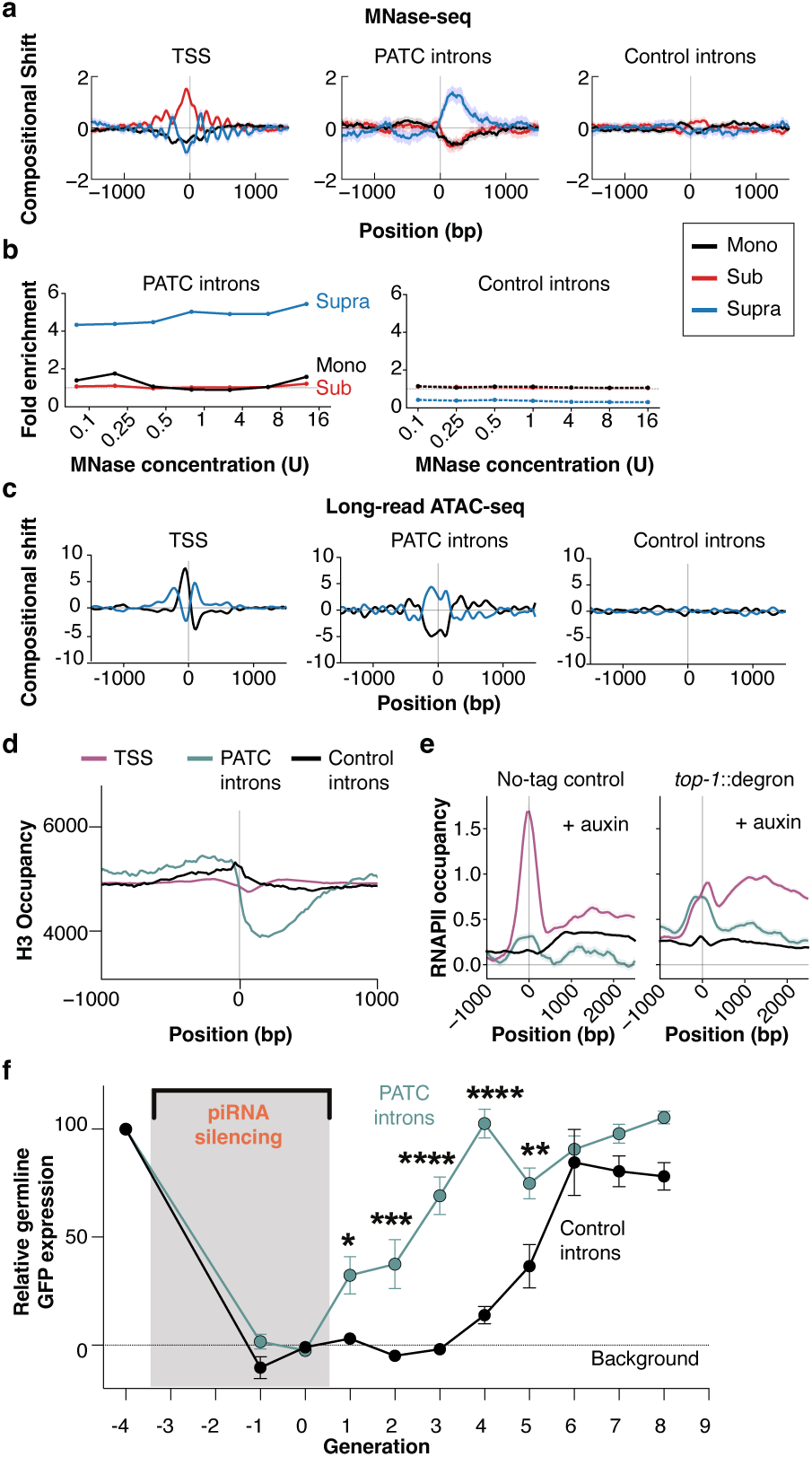
PATCs form topological nucleosome-depleted barriers. **a**, Previously published MNase-seq [41] compositional shift centered on TSS, PATC-rich introns, and matched control introns. Fragment classes: sub-nucleosomal (< 120 bp, red), mononucleosomal (120-165-bp, black), and supranucleosomal (> 165 bp, blue). **b**, Fold enrichment of recovered fragments over PATC-rich (solid lines, top) and matched control introns (dashed line, bottom) versus MNase concentration. **c**, Long-read ATAC-seq compositional shift for mono-(150-260-bp) and supra-nucleosomal (> 300-bp) fragments. **d**, Previously published histone 3 (H3) CUT&RUN signal [42] centered on TSS (pink), PATC-rich introns (teal), and matched control introns (black). **e**, Previously published RNAPII ChIP-seq profiles [38] for a no-tag control and a topoisomerase-depleted *top-1* ::*degron* strain in the presence of auxin. **f**, Quantification of GFP fluorescence recovery from piRNA silencing for matched single-copy P*mex-5* ::*gfp* reporters (PATC-rich versus control introns) inserted in permissive chromatin (*ttTi5605* II insertion site). Mean GFP fluorescence ± SEM (PATC introns: N = 5, Control introns: N = 4 strains per condition, 13-18 animals imaged per generation). P values: Gen 1: 0.0389, Gen 2: 0.0005, Gen 3: <0.0001, Gen 4: <0.0001, Gen 5: 0.0021, two-way ANOVA with Bonferroni multiple comparison adjustment.

If this structure is indeed topological, it should impede transcription, particularly when torsional stress cannot be relaxed. We analyzed previously published RNAPII ChIP-seq data following the acute depletion of topoisomerase I (TOP-1) [38]. While TOP-1 depletion generally increases polymerase density in gene bodies, this stalling is amplified over PATC-rich introns compared to controls (Fig. 5e). This further suggests that PATCs impose a physical barrier that interacts with the topological state of the chromatin.

### Resilience to silencing ensures homeostasis

In *C. elegans*, small RNAs initiate heritable silencing in the germline by destabilizing transcripts and directing the deposition of repressive histone marks [43]. We reasoned that if PATCs disrupt canonical nucleosome organization, they may not block the initial cytoplasmic recognition and destruction of a target transcript, but instead interfere with the downstream HRDE-1-dependent coupling between small-RNA targeting and chromatin-based inheritance [44].

To distinguish these possibilities, we targeted matched single-copy reporters—differing only by the presence or absence of PATC-rich introns—with synthetic piRNAs to induce acute germline silencing [45]. Both reporters were effectively silenced, establishing a uniform repressed baseline (Fig. 5f). However, the subsequent dynamics differed strikingly. While control reporters remained fully silenced for three generations after the trigger was removed, the PATC-containing reporter displayed immediate resilience: fluorescence levels recovered to ∼ 30% in the very first generation, with full recovery within four generations (Fig. 5f). Thus, PATCs limit the persistence of small-RNA-induced silencing across generations. This supports a model in which the disruption of canonical nucleosome organization weakens the establishment or maintenance of repressive chromatin memory, allowing a rapid return to transcriptional activity (Supplementary Fig. S9).

### Convergent evolution of sequence-encoded mechanics

While PATCs represent an evolutionary extreme of transposon co-option, we hypothesized that the underlying principle—localized minima in DNA bending energy—might be a shared functional feature of metazoan genomes. To test this, we deployed our TBE model across the animal kingdom, leveraging its unique ability to scale biophysical predictions to entire genomes. This broad survey reveals that such “high-pinning” sequences are ubiquitous, though the degree of genomic amplification varies (Fig. 6a). In the Pacific oyster (*Magallana gigas*), for example, independent expansions of Helitron-derived satellite DNA [46] have generated distinct genomic fractions that resemble PATCs in both sequence periodicity and physical character (Supplementary Fig. S10). This suggests that Helitrons can act as recurrent evolutionary vehicles for DNA sequences with these mechanical properties, expanding their known role in mobilizing regulatory elements [47].

**Figure 6:**
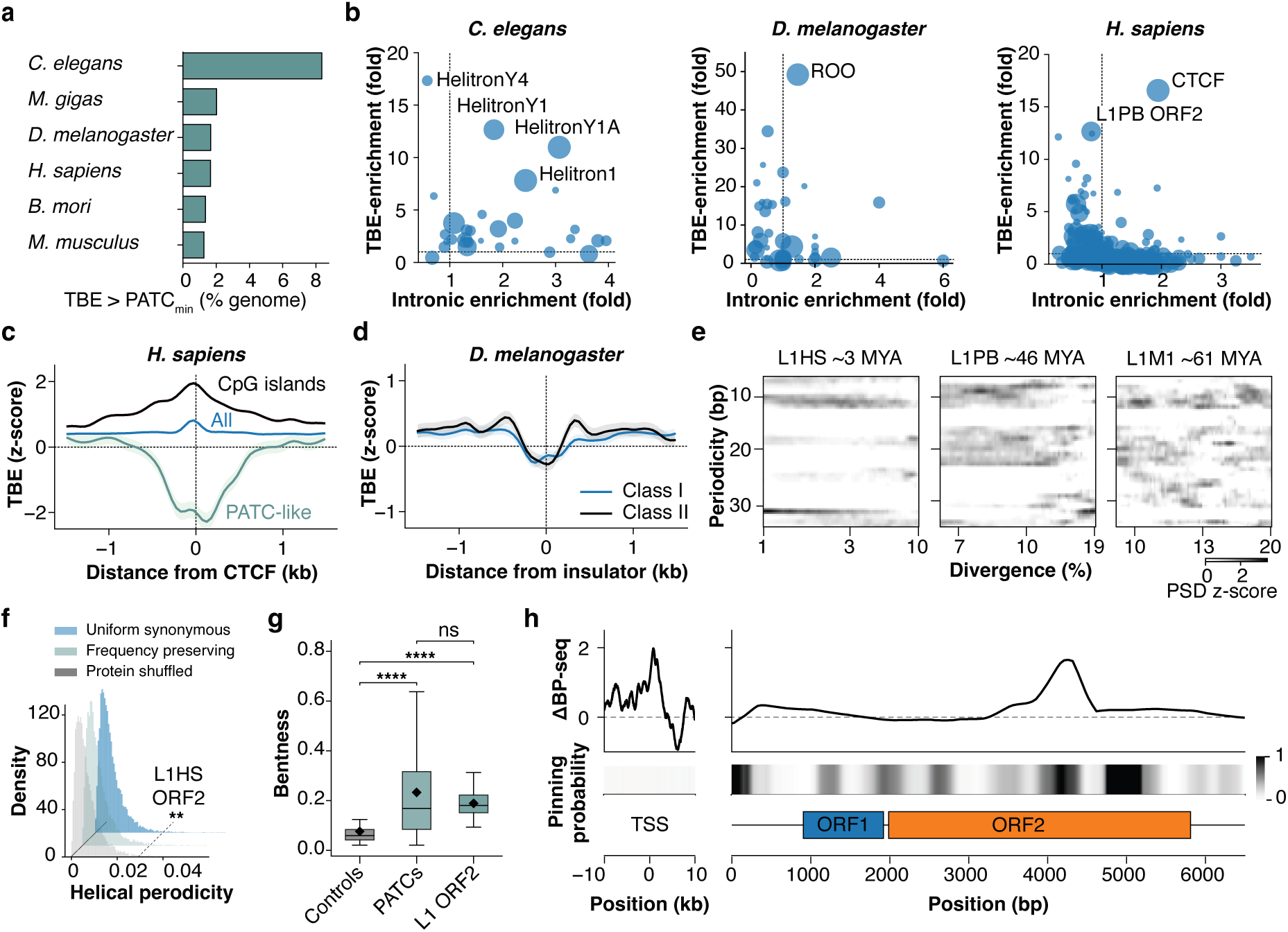
Evolutionary convergence of sequence-encoded mechanics. **a**, Genomic distribution of high-pinning sequences. Fraction of metazoan genomes containing sequences with TBE values within the range of *C. elegans* PATCs. **b**, Enrichment of pinning elements at functional loci. Analysis of the PATC-like pinning potential at annotated repeats and insulator binding sites relative to their intronic enrichment in *C. elegans, H. sapiens, and D. melanogaster*. (**C-D**) Bending energy profiles at canonical insulators. Predicted TBE (z-score) centered on experimentally defined human CTCF sites (C) and *Drosophila* Class I and Class II insulator sites (D) [82]. **e**, Evolutionary decay of L1 periodicity. Spectrogram analysis of L1 subfamilies of varying ages (MYA: million years ago). Note the robust ∼10-bp helical periodicity in young elements (L1HS) and its progressive erosion in older lineages (L1M1). **f**, Synonymous simulated recoding of L1HS ORF2. Distribution of helical periodicity scores for *in silico* recoded L1HS ORF2 sequences. The native sequence (dashed line) is significantly more periodic than all null models (One-sided empirical randomization tests (n = 10,000 recoded sequences per null model; +1 correction): uniform synonymous, P = 0.00690; codon frequency preserving, P = 0.00640; protein shuffled, P = 0.00760.) including those preserving L1 codon frequencies. **g**, Structural quantification. Comparison of sequence-encoded bentness for *C. elegans* PATCs, control introns, and the human L1 ORF (Two-sided Mann–Whitney U tests (n = 100 sequences per group): control versus PATC, U = 1,667, P = 3.87 × 10*^−^*^16^; control versus L1 ORF2, U = 710, P = 1.06 × 10*^−^*^25^; PATC versus L1 ORF2, U = 4,802, P = 0.629. ****P < 0.0001; ns, P ≥ 0.05.). **h**, *In vivo* validation of L1 topological barriers. Aggregate normalized pinning probability (bottom) and mean previously published BP-seq signal [52] (top) aligned to the intronic L1 consensus sequence and human Transcription Start Sites (TSS).

To compare genomes with distinct energetic baselines, we developed a genome-normalized pinning probability, defined relative to the median topological DNA bending energy of each host species. Given that *C. elegans* PATCs evolved within introns, we reasoned that analogous mechanical elements in other species might—like their nematode counterparts—accumulate within transcribed regions or regulatory boundary elements. We therefore screened annotated repeats and insulator binding sites, plotting their predicted pinning strength against their intronic enrichment (Fig. 6b). In *C. elegans*, the intersection of these mechanical and transcriptional signatures specifically identifies the expected Helitron families, validating that our analytical framework captures *bonafide* pinning elements. Strikingly, extending this analysis to the human genome identifies CTCF binding sites [48] as the primary outliers; these sites segregate into a flexible “mechanical” class that is distinct from the rigid CpG-rich boundaries previously described [49, 50] (Fig. 6c). This mechanical signature extends to *Drosophila*, where Class I and Class II insulators similarly map to deep TBE minima (Fig. 6d), suggesting that supercoiling elements are a convergent feature of boundary elements across phyla. Our theoretical predictions are corroborated by recent high-throughput Loop-seq assays, which demonstrate that CTCF sites possess extreme, sequence-encoded intrinsic cyclizability that extends well beyond the canonical binding motifs [51]. Furthermore, sequence-based models trained on these empirical cyclizability measurements independently predict the same extreme bendability for the *C. elegans mrpl-40* intron (Fragment Ω4)—a prototypical PATC sequence [51]. The convergence of these two distinct approaches—empirically trained models and our analytical, physics-based TBE calculations— provides robust orthogonal validation of a shared mechanical code.

Beyond CTCF, the most distinct human signal maps to the L1 ORF2 coding region, which is strongly enriched for PATC-like DNA bending topology (Fig. 6b). Reversing the evolutionary trajectory observed for nematode Helitrons (Fig. 1e), spectrogram analysis of L1 subfamilies reveals that robust 10-bp helical periodicity is prominent in the youngest elements (L1HS-PA, currently active to ∼3 MYA) but progressively erodes in older, divergent lineages (Fig. 6e). This periodic signature is dual-encoded within the ORF2 protein sequence via specific positional codon usage. *In silico* recoding of ORF2 demonstrates that this helical periodicity is exceptionally constrained; all synonymous null models—including those preserving L1 codon frequencies—completely abolish the mechanical signal (P < 0.01), Fig. 6f). Similar to *C. elegans* PATCs, this periodicity encodes intrinsic curvature (Fig. 6g) that tracks to localized topological stress *in vivo*. Pinning predictions align with independent, previously published experimental BP-seq measurements [52], identifying topological stress within active L1 ORF2 families (Fig. 6h). Furthermore, L1 elements exhibit a PATC-like “supra-nucleosomal” ATAC-seq [53] footprint, a feature absent in the extinct LINE-2 family (Supplementary Fig. S11).

ORF2 is known to impede its own transcription through sequence-intrinsic properties—a form of self-attenuation proposed to limit the fitness cost of unchecked amplification [54]. Our findings suggest a possible physical basis for this phenomenon: the ORF2 coding sequence encodes intrinsic curvature and high topological pinning potential, features that could stall RNA transcription. More speculatively, this same structural feature could help active L1 elements navigate the conflict between expression and host repression: like Helitron-derived PATCs in *C. elegans*, L1 ORF2 may exploit sequence-encoded mechanics to reduce the probability that transcriptionally active elements are converted into stably silenced chromatin. Thus, PATCs and L1 ORF2 may represent opposite evolutionary uses of a similar physical principle: host genomes can co-opt transposon fossils as protective chromatin elements, whereas active transposons may retain related mechanical features to modulate their own expression and persistence.

## Discussion

Together, these findings suggest that genomic insulation is not merely a function of protein binding, but can also emerge from physical properties encoded directly within the DNA polymer. In *C. elegans*, Helitron-derived minisatellites evolved a 10-bp periodicity that creates intrinsic curvature, lowers the energetic cost of local DNA deformation, disrupts canonical nucleosome organization, and protects active germline loci from progressive and heritable silencing. This provides a mechanistic explanation for how a genome lacking canonical CTCF-like insulators can nevertheless maintain functional separation between active and repressive chromatin domains.

A key distinction is that PATCs do not resemble classical focal insulators. They are long, intronic, and often distributed across kilobases within active genes. Their boundary activity is therefore unlikely to arise from a single protein-binding event at a precise domain border. Instead, PATCs appear to form extended active-side barrier zones: regions within transcribed genes that locally alter DNA/chromatin structure and resist the encroachment or stabilization of repressive chromatin. This distributed nature explains why PATCs are enriched on the active side of chromatin-state transitions rather than being centered at boundaries.

We propose that the most parsimonious physical model for this behavior is the formation of localized, supercoiling-responsive DNA structures, potentially plectoneme-like conformations. Under this model, PATCs act as energetic sinks for transcription-induced torsional stress. Their phased A/T tracts lower the cost of bending the DNA, increasing the probability that supercoiling-induced deformation is absorbed locally. Such a structure would be expected to compact the DNA, exclude or destabilize canonical nucleosomes, and create a discontinuity in the chromatin fiber. This provides a simple physical route by which an intronic sequence could behave as a barrier: not by recruiting a classical insulator protein, but by locally altering the substrate through which repressive chromatin must propagate.

Importantly, this model does not require PATCs to act in a protein-free manner. The digestion-resistant, nucleosome-depleted footprint of PATC-rich introns could reflect intrinsic DNA folding, torsionally constrained DNA, associated protein factors, or a combination of these mechanisms. These possibilities are not mutually exclusive. A sequence-encoded DNA structure could recruit, exclude, or reposition proteins; conversely, protein binding could stabilize a structure initially biased by DNA mechanics. Thus, intrinsic DNA mechanics and trans-acting chromatin factors should be viewed as coupled layers rather than competing explanations.

This model also connects PATCs to transgenerational epigenetic inheritance. In the *C. elegans* germline, small-RNA pathways can convert transient transcript targeting into stable chromatin memory. For such memory to persist, repression must be coupled back to the chromatin template, likely through Argonaute binding to nascent transcripts in the nucleus and local deposition on nearby nucleosomes [44]. PATC-rich regions may disrupt this transgenerational inheritance mechanism. Their nucleosome-depleted yet digestion-resistant footprint, together with their rapid recovery from piRNA-induced silencing, suggests that PATCs do not block post-transcriptional small-RNA mediated transcript silencing, but limit the ability of silencing signals to stabilize into nuclear inherited chromatin states. In this sense, PATCs may act as mechanical breaks in the epigenetic memory circuit to prevent inadvertent silencing of endogenous genes by the strong repressive environment in the germline [12].

The evolutionary origin of PATCs makes this mechanism especially striking. Transposable elements are usually viewed as targets of germline repression, yet in this case ancient Helitron fossils appear to have been remodeled into sequences that protect host genes from that same repressive machinery. This expands the concept of transposon co-option. While TE-derived insulators have often been understood as dispersed protein-binding motifs [55–57], PATCs suggest a different route: the co-option of repetitive DNA as physical material. Helitrons were not simply carrying regulatory motifs; their internal minisatellites provided an evolvable substrate from which distributed mechanical barriers could be written. Repeated monomers are especially suited to this process because weak sequence-level bending biases can be amplified across many copies, allowing small mutational changes to generate extended mechanical domains.

The broader comparative analyses suggest that this principle is not unique to nematodes. Human CTCF sites, *Drosophila* insulators, oyster Helitron-derived satellites, and human L1 ORF2 sequences all show signatures of sequence-encoded bendability or topological pinning potential. These elements do not share a common sequence origin, and they need not form identical structures. What appears shared is a physical tendency: the ability of primary sequence to bias local DNA deformation under torsional stress. Thus, the convergence lies not in a conserved motif, but in a shared mechanical solution [51].

The emergence of CTCF is closely associated with the radiation of bilaterians [9], yet the need to partition active and repressed chromatin is likely as ancient as eukaryotic chromatin itself. Sequence-encoded mechanics may therefore represent an older substrate for insulation, onto which protein-based boundary systems were later layered. In this view, canonical insulator proteins may not create boundaries on a homogeneous polymer. Instead, they may recognize, stabilize, or regulate a pre-existing mechanical landscape encoded in DNA. CTCF, cohesin, topoisomerases, chromatin remodelers, and small-RNA pathways may all act on this landscape, but the landscape itself is partly written by sequence. *C. elegans* provides an unusually clear example because the mechanical component became amplified genome-wide through Helitron expansion.

Several questions remain open. Our data support sequence-encoded curvature, topological responsiveness, nucleosome disruption, and resistance to heritable silencing, but they do not uniquely define the precise in vivo structure formed by PATCs. Plectoneme-like conformations [28] provide one coherent physical model, but direct structural experiments will be required to test this. Acute TOP-1 perturbation followed by ATAC-seq or MNase-seq could determine whether the PATC-associated protected-fragment signature is enhanced under torsional stress. Assaying acute and inherited silencing in HRDE-1 or nuclear RNAi mutants could determine whether PATCs specifically limit chromatin-based inheritance rather than cytoplasmic silencing. Finally, reconstituted nucleosome arrays containing PATC-rich sequences under supercoiling stress would provide a direct biophysical test of whether these elements disrupt nucleosome organization through DNA mechanics.

We therefore propose that sequence-encoded DNA mechanics represent an ancient and evolutionarily accessible substrate for chromatin insulation. In host genomes, fossilized transposable elements can be repurposed as protective barriers that preserve gene expression in repressive environments. In active selfish elements such as L1, related mechanical features may instead tune transcriptional output and help evade host repression. The same physical principle may therefore be harnessed by both sides of the genome conflict: by hosts to protect endogenous genes, and by mobile elements to persist within the chromatin environments that seek to silence them.

## Methods

### Genomic Data and Annotations

The *C. elegans* N2 reference genome (version WS290) and the human reference genome (GRCh38/hg38) were used for all analyses. Gene annotations were obtained from WormBase. Repetitive element annotations, including Helitron and LINE families, were derived from the Dfam database [20] utilizing consensus sequences defined by Repbase [58]. Family names were shortened for textual clarity. For population genomics, genome assemblies of 17 wild *C. elegans* isolates were obtained from previously published sources [23–25].

### PATC Identification and Quantification

Periodic A/T Cluster (PATC) scores were calculated on a per-base level using the algorithm described by Fire *et al*. [14]. A phasing threshold of 90% was applied for all analyses. PATC density for a given region was defined as the average per-base PATC score over the length of that region.

### Co-localization Analysis

To quantify the contribution of repeat families to the genomic PATC signal, we intersected genomic coordinates of PATCs with repeat annotations. The fraction of the total PATC signal that co-localized with each repeat family was then calculated. The statistical significance of this co-localization was determined through a permutation test (N=10,000), where repeat annotations were randomly shuffled across the genome to establish a null distribution.

### Identification and Dating of Minisatellites

To identify minisatellites, we first ran Tandem Repeats Finder [59] on Helitron consensus sequences using parameters (2 7 7 80 10 60 1000). Redundant or overlapping annotations (> 25% overlap) were resolved by selecting the representative with the shortest consensus sequence. Minisatellites were clustered by sequence similarity using an edit distance threshold of < 30%. Clustered sequences were aligned using MUSCLE [60] to produce the cluster consensus sequence.

### Mapping PATC Density onto Consensus Sequences

To generate the PATC density map for each Helitron family (Fig. 1b), we mapped the PATC signal from all individual genomic insertions to their respective consensus sequences. Per-base PATC scores were extracted for the genomic coordinates of every annotated insertion within a family. To account for length variations driven by indels, the PATC signal from each insertion was rescaled to the consensus coordinate system via linear interpolation. These rescaled profiles were then aggregated to produce a single, aggregate PATC density profile for each Helitron consensus sequence.

### HMM-based Identification and Dating of Minisatellites

To annotate Helitron-derived satDNA, we built a sensitive profile Hidden Markov Model [61] for the PATC-precursor minisatellite. Initial seed sequences were identified by finding high-density genomic clusters of the 35-bp monomer using stringdecomposer [62]. These sequences were dereplicated using cd-hit-est [63]. Aligned sequences (mafft) [64] were used to construct the profile HMM via hmmbuild. We then used this HMM to perform a genome-wide search with nhmmer [61]. Loci < 200 bp or overlapping other transposable element families were excluded. The evolutionary age was estimated by nucleotide divergence from the HMM consensus sequence. Elements were classified as ‘Young’ (< 20%), ‘Intermediate’ (20–30%), and ‘Ancient’ (> 30%). Divergence times (MY) were estimated using established mutation rates and generation times [65].

### Gene Expression and Genomic Context Analysis

To analyze the relationship between minisatellite evolution and genomic context, we associated each curated minisatellite instance with previously published tissue-specific gene expression data [66]. This allowed us to assign a tissue expression pattern (e.g., ‘Germline’, ‘Ubiquitous’, ‘Neurons’) to each element based on its host gene. Minisatellites not overlapping annotated gene bodies were classified as ‘intergenic’. Tissue-specific categories were standardized by merging closely related subgroups (e.g., ‘Ubiquitous’ and ‘Ubiquitous-Biased’ into ‘Ubiquitous’) to ensure robust statistical comparisons.

### Power Spectral Density Analysis of Periodicity

To analyze the evolution of periodicity in minisatellites and L1 sequences, we performed Power Spectral Density (PSD) analysis. Minisatellite sequences were sorted by evolutionary age (TN93 divergence) and grouped into bins. For each sequence, a binary signal was generated by marking the start positions of A/T-tracts (defined as ≥ 4 consecutive A or T bases) with a ‘1’. The power spectrum for these binary signals was computed using the scipy.periodogram [67] function. Resulting spectra were averaged across all sequences within each evolutionary age bin. To highlight age-dependent shifts in periodicity, the mean PSD vector for each bin was Z-score normalized relative to the other bins. The final matrix, representing the normalized power of each periodicity at each evolutionary age, was visualized as a heatmap.

### Monomer-Folding Alignment Method

To analyze position-specific mutation rates, we developed a “monomer-folding” strategy to map all genomic minisatellite sequences to a common 35 bp monomer coordinate system. First, all curated genomic instances of the minisatellite were aligned to the full-length HMM consensus sequence using hmmalign. We then used stringdecomposer [62] to identify the precise locations of the 35 bp monomers within the full-length consensus. A base-level map was created by performing a semi-global alignment of each of these monomer instances against the canonical 35 bp monomer sequence. This produced a final mapping where every position in the full-length consensus was assigned a corresponding coordinate on the 35 bp monomer, allowing for the direct comparison of mutations across all genomic copies.

### Mutability Rate Calculation and ssDNA Structure

Aligned sequences were partitioned into ‘Young’, ‘AncientLowPATCs’, and ‘AncientHighPATCs’. For each position in the 35 bp monomer, we calculated a mutability rate as the total number of observed mutations divided by the total number of observed bases. Excess mutability was calculated as:

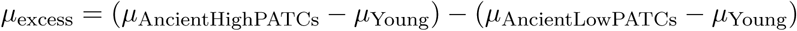

This isolates the mutational signature specifically associated with PATC evolution by subtracting the background drift found in low PATC ancient elements. Separately, to test a structure-driven mutational mechanism, we predicted the secondary structure of the 35-bp monomer. We used RNAfold [68] with DNA parameters to calculate the pairing probability for each base. The probability of a base being in a single-stranded DNA loop was determined by (1 − ∑ *P*_pairing_), which was correlated with the excess mutability.

### Forward Evolutionary Simulation

The simulation was initialized with the ancestral HMM consensus sequence. At each simulated generation, every base in the sequence was subjected to stochastic mutation based on the position-specific mutability rate corresponding to its location on the 35-bp monomer. We performed 500 independent replicate simulations for three models:(i) High PATC Model: mutability rates derived from the ‘AncientHighPATCs’ group; (ii) Low PATC Model: mutability rates derived from the ‘AncientLowPATCs’ group; and (iii) Shuffled Control: high PATC mutability rates applied to a shuffled ancestral sequence.

### Population Genomics Analysis

To analyze the conservation and evolution of Helitron-derived minisatellites across wild populations, we identified orthologous loci in 17 *C. elegans* wild isolate genomes. For each minisatellite locus in the N2 reference, we extracted 10 kb of flanking sequence on both sides. These flanks were used as queries to search each wild isolate genome using lastal [69]. A locus was considered orthologous if both flanks mapped uniquely to the same contig, in the correct orientation and strand, with alignment ends located within 1500 bp of the predicted insertion site. Based on the distance between the mapped flanks, each locus was classified as: (i) Conserved: insert size was within 10% of the N2 reference; (ii) Partially Conserved: insert was present but the size had diverged by more than 10%; (iii) Absent: insert size was less than 10 bp. Evolutionary shifts in PATC density (’Increased’, ‘No Change’, or ‘Decreased’) were determined by comparing scores between wild isolates and the N2 reference.. Finally, we used a Chi-squared test to determine if the direction of change was significantly associated with the host gene’s expression context.

### Topological Bending Energy (TBE) Model

We developed an algorithm to calculate the mechanical energy required to deform a DNA segment into the tight loop geometry characteristic of a plectoneme tip. The TBE model assumes a 73-bp window and a 240*^◦^* bend angle based on established plectoneme geometries [28]). The underlying mechanical properties were modeled using an Ising model [33] parameterized from large-scale molecular dynamics simulations, treating the DNA polymer as a chain of interacting steps. For a given input sequence, we computed the equilibrium geometric parameters (µ⃗) and 12×12 stiffness matrices (***K***). Relaxed conformations and effective compliance matrices (J) were derived by inverting (K) and solving for the energy minimum of the unconstrained polymer. Plectoneme nucleation energy (ΔF) was defined as the cost to deform the 73-bp window into a loop with a total bend magnitude of 240*^◦^*. To compare pinning potential across genomes with distinct baselines (ΔF) values were converted into calibrated pinning probabilities (P*_pin_*). We modeled the probability of a specific site capturing supercoiling using a Boltzmann weight relative to the genome-wide median bending energy (E*_ref_*):

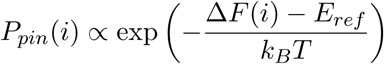

The proportionality constant was calibrated such that the probability density matches a target expectation (e.g., P ≈ 10*^−^*^6^ at the median energy) to ensure that pinning sites are identified based on their relative potential within a specific genomic context.

### Identification of Chromatin Boundaries

To identify robust transitions between active (H3K36me3) and repressive (H3K27me3) chromatin domains, we developed a custom boundary-calling algorithm. We first generated a composite signed chromatin track defined as the log _2_-ratio of H3K36me3 to H3K27me3 enrichment, where positive values denote active chromatin and negative values denote repressive chromatin. Boundaries were defined as genomic loci exhibiting a sustained transition between these states. To minimize false discoveries driven by local noise, we imposed a strict purity threshold. A window was considered to “pass” only if the fraction of bases matching the expected sign exceeded 0.9. The boundary coordinate was defined as the center of this transition zone. We employed an asymmetric window size (L*_pos_* = 2.5 kb, L*_neg_* = 10 kb as a minimum). This asymmetry accounts for the observation that H3K27me3 forms broad domains, unlike H3K36me3, which is often restricted to single gene bodies [10].

### Hi-C Data Analysis

Processed Hi-C contact maps (in .hic format) were obtained from previously published datasets [38, 39]. To visualize boundaries, we performed a metaplot (pileup) analysis using hicstraw [70] and custom scripts. For each boundary, we extracted a local interaction matrix centered on the boundary coordinate. We utilized a window size of ± 10 kb at 500-bp resolution. To correct for genomic distance biases and sequencing depth, we used Knight-Ruiz (KR) normalized Observed/Expected (O/E) values. These matrices were averaged element-wise to generate a global “pileup” map.

### BP-seq, MNase-seq and ATAC-seq Fragment Size Analysis

To dissect the chromatin landscape based on fragment size, paired-end MNase-seq reads were stratified into three distinct classes: sub-nucleosomal (< 120 bp), mono-nucleosomal (120 − 165 bp), and supra-nucleosomal (> 165 bp). Similarly, ATAC-seq reads were partitioned into subnucleosomal (< 120 bp), mono-nucleosomal (170 − 230 bp), and supra-nucleosomal (> 300 bp). These size thresholds reflect the distinct biochemical properties of each enzyme: MNase digests linker DNA, whereas Tn5 transposase inserts into it [71, 72]. Alignment files were converted into genome-wide coverage tracks and normalized by Counts Per Million (CPM). Relative enrichments were visualized using log2-ratio tracks (e.g., log_2_(Supra/Mono)). For BP-seq, log_2_(treated/gDNA) tracks were generated as described previously for *C. elegans* [36]. For human cell line data, previously processed ChBPseq bigwig files were utilized. All reads were aligned to the *C. elegans* N2 reference genome using Bowtie2 [73] in –local –very-sensitive-local mode.

### In Silico Recoding and Helical Periodicity Analysis of L1 ORF2

Pairwise distances between AT-tracts (≥ 4 bp) were used to construct spacing histograms. To quantify periodicity, we applied a Fast Fourier Transform (FFT) to obtain the power spectral density. A helical periodicity score was defined as the ratio of the peak FFT power within the helical band (10-11 bp) to the total spectral power. To determine whether the observed 10-bp helical periodicity is a highly constrained feature of positional codon usage, we compared the native ORF2 sequence against 10,000 permutations of three null models: (i) Uniform Synonymous: native amino acid sequence was fixed with synonymous codons selected uniformly at random; (ii) Frequency Preserving: native amino acid sequence was fixed, and native L1 codons were permuted among matching amino acids, preserving L1 codon frequencies; (iii) Protein Shuffled: amino acid order was randomized while maintaining the overall codon usage. Empirical P-values were calculated by comparing the native ORF2 helical periodicity score to the distributions generated by the three null models.

### AFM imaging in liquid

Ten microliters of sample were deposited onto a freshly cleaved mica surface mounted in a liquid cell. After a 10-minute incubation, the liquid cell was filled with 0.5 mL of 0.5x TBE buffer supplemented with 200 mM MgAc_2_ and 50–100 mM NiCl_2_. AFM measurements were carried out using a Bruker Dimension Icon AFM operated in ScanAsyst mode and equipped with a SCANASYST-FLUID+ probe (Bruker). Cantilevers with a resonance frequency of 100– 200 kHz and a nominal spring constant of 0.7 N m*^−^*^1^ were used. Images were acquired at a resolution of 256 × 256 pixels. A low imaging force (∼200 pN) and a fast scan rate of 2-3 Hz were applied to minimize sample deformation.

### Dry-state AFM imaging

Ten microliters of the DNA sample (3 nM) were deposited onto mica and incubated for 10 minutes. The surface was rinsed three times with 200 µL of ultrapure water and dried overnight in a desiccator. AFM measurements were performed in intermittent contact mode with an FESPA-V2 probe (Bruker). Cantilevers with a resonance frequency of 50-100 kHz and a spring constant of 2.8 N m*^−^*^1^ were used. Images were acquired at a resolution of 256 × 256 pixels and processed using Gwyddion software.

### Total Curvature from AFM Images

AFM images of plasmid DNA were processed to extract a one-pixel-wide backbone trace for each molecule. DNA features were isolated by thresholding to generate a binary mask, followed by skeletonization to obtain a centerline representation. The resulting skeleton was converted into an ordered polyline to produce a continuous backbone trace. Arc length (s) along the trace was computed cumulatively from successive points. Local curvature k(s) was estimated using finite-difference derivatives of the tangent direction along the smoothed centerline. Total curvature was quantified as the total absolute curvature (T AC*_abs_*) accumulated along the trace, where L is the total trace length:

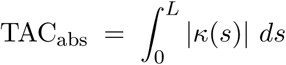

### Worm Strains and Maintenance

*Caenorhabditis elegans* strains were maintained using standard methods [74]. For routine maintenance, transgenesis, and injections, worms were cultured on Nematode Growth Medium (NGM) plates seeded with *E. coli* OP50. For large-scale experiments requiring high biomass, such as ATAC-seq, worms were grown on large NGM plates seeded with *E. coli* NA22. All strains were cultured at 20–25*^◦^*C. A complete list of strains, including specific genotypes, is provided in **Supplementary Table 1**. All experiments were performed in accordance with KAUST Institutional Biosafety and BioEthics Committee (IBEC), approval no. 17IBEC34.

### Transgenesis

Following established direct-insertion MosSCI protocols [16, 75], we generated single-copy transgene insertions by microinjecting DNA into the germline of *C. elegans*. Selection for successful integrations was provided by the *cbr-unc-119(+)* rescue marker in an *unc-119(ed3)* mutant background. We identified stable, homozygous transgenic lines based on Unc-119 rescue and the absence of extrachromosomal arrays marked by mCherry co-injection markers. Transgenes were targeted to two distinct genomic environments: the germline-permissive *ttTi5605* site on Chromosome II [75] or the repressive *oxTi173* universal MosSCI site on Chromosome V [15]. Following successful identification of the integrated lines, all strains were maintained continuously at 25*^◦^*C to prevent the onset of progressive transgene silencing [11]. A complete list of strains, including their specific insertion sites and associated genotypes, is provided in **Supplementary Table 1**.

### Molecular Biology and Transgene Assembly

#### Transgene Design and Synthesis

Transgenes were optimized and assembled as described in Aljohani *et al.* [16]. The green fluorescent protein (*gfp*) coding sequence was codon-optimized for *C. elegans* to maximize expression by selecting highly expressed codons. To minimize potential germline silencing, the sequence was depleted of piRNA-binding sites [17], and internal BsaI restriction sites were removed. Because long endogenous introns cannot be reliably generated via direct gene synthesis, an initial *gfp* variant was synthesized by Twist Bioscience. This variant contained one short endogenous intron and three additional synthetic introns carrying unique Golden Gate acceptor sites to facilitate subsequent modifications.

#### Insertion and Assembly

This sequence-verified *gfp* served as an “acceptor” transgene for the insertion of various long endogenous introns. Using the NEBridge Golden Gate Assembly Kit (BsaI-HF v2, NEB cat. no. E1601L), the three synthetic introns were exchanged in a one-pot reaction for endogenous *C. elegans* introns of various lengths. These introns—which included PATC-rich sequences, matched non-PATC-rich controls, or young Helitron-derived elements— were individually amplified and sequence-verified prior to insertion into the *gfp* backbone. The resulting plasmids were validated by restriction digestion to confirm correct assembly.

#### Expression Vector Assembly

Final expression constructs were generated via Gateway LR cloning (Gateway LR Clonase II Enzyme mix, ThermoFisher cat. no. 11791020) according to the manufacturer’s instructions. Transgenes were moved into destination vectors designed for use with universal MosSCI single-copy insertion sites, containing the *cbr-unc-119* rescue marker [76] and left/right genomic homology regions. This design allowed for the targeted insertion of transgenes at various genomic locations, including the centers and arms of autosomes, as detailed for each strain in **Supplementary Table 1**. The final expression vectors were validated by restriction digestion and gel electrophoresis. A comprehensive list of all plasmids, including full DNA sequences, is provided in **Supplementary Table 2**.

#### Data Availability of Sequences

To facilitate the reuse and inspection of the transgenes described, annotated GenBank files containing complete feature maps (including codon-optimized exons, specific intron coordinates, and regulatory elements) are provided as **Supplementary Data 1**. These files correspond to the plasmids listed in **Supplementary Table 2**. All sequences were managed and annotated using ApE (A plasmid Editor) [77].

### Germline Fluorescence Quantification

To standardize the timing of transgene analysis, P_0_ animals were injected and placed individually on NGM plates at 25*^◦^*C. Populations were allowed to propagate until starvation, and stable single-copy insertions were identified in the F_2_ generation. From each successful injection, a single F_2_ adult was picked to establish an independent clonal line. To assess the initial frequency of germline expression, F_3_ progeny were utilized for all primary quantifications. The integrity of each insertion was verified by monitoring constitutive somatic GFP expression driven by the *eft-3* promoter, which served as an internal control for the presence of the transgene. For experiments monitoring the stability of expression over time (Figure 4E), animals with more than 80% germline expression from various control (pCFJ2407, pCFJ2408, pCFJ2409, pCFJ2410) and PATC (pCFJ2371, pCFJ2373, pCFJ2375, pCFJ2405, pCFJ2412, pCFJ2413) plasmids were maintained at 25*^◦^*C and re-evaluated in the F_5_ generation. Imaging and scoring was performed on worms anesthetized with 50 mM sodium azide and mounted on 2% agarose pads for visualization. Imaging was performed using a 42× oil immersion objective on a compound microscope. Due to the stochastic nature of transgene silencing [15], germline expression was scored as a binary (“on” or “off”) trait in at least 11 independent animals for each independent insertion line. A line was considered “expressing” only if GFP fluorescence was visible in the germline syncytium or oocytes, independent of somatic expression levels.

### Transgenerational Silencing Recovery Assay

#### Strain Establishment and Validation

Single-copy transgene insertions were generated via MosSCI as previously described, utilizing the germline-specific *mex-5* promoter [78] to drive *gfp* variants containing either (i) 250-bp control introns (pMNK19) or (ii) 250-bp PATC-rich introns (pMNK20) at 25*^◦^*C. Two representative non-silencing strains, CFJ22 (control) and CFJ23 (PATC-rich), were established and validated to ensure that initial germline fluorescence levels were statistically indistinguishable. The susceptibility of these transgenes to small-RNA-mediated repression was independently verified using piRNAi [45] and RNAi feeding [79] against the codon-optimized *ce-gfp* sequence.

#### Silencing and Recovery Protocol

To initiate transgenerational silencing, the T59 piRNAi fragment—encoding six synthetic piRNAs directed against *gfp*—was injected into young adult animals, designated as generation “−4”. From these injections, we established five independent control lines (CFJ22 derivatives) and four PATC-rich lines (CFJ23 derivatives) that maintained stable extrachromosomal arrays capable of silencing the integrated transgene. Transgenic populations were monitored and imaged at generations “−1” and “0” to confirm the silenced state. The silencing stimulus was subsequently removed by selecting progeny that had lost the extra-chromosomal piRNAi array. The recovery of germline fluorescence was subsequently quantified across generations “1” through “8”.

#### Fluorescence Quantification

Young adult transgenic animals were anesthetized with 10 mM sodium azide and mounted on 2% agarose pads for visualization. Imaging was performed on a compound microscope utilizing a 42x× oil immersion objective, with 13–18 independent animals imaged for each generation and genotype (five control and four PATC strains). Non-transgenic animals were imaged under identical settings to determine the baseline autofluorescence in the GFP channel. Fluorescence intensity was quantified using a standardized ImageJ script, and the mean background-subtracted signal was recorded for each animal [80]. All transgenes are listed in **Supplementary Table 2**.

### Long-read ATAC-seq

#### Growth and Synchronization

To obtain sufficient biomass, unsynchronized populations were expanded on large NGM plates (100 × 15 mm) seeded with NA22. Gravid adults were harvested and subjected to bleaching (16 mL sodium hypochlorite, 20 mL 10 N NaOH, 64 mL H_2_O) to release embryos. Embryos were incubated overnight in M9 buffer on a rotator to synchronize at the L1 stage, followed by two additional rounds of expansion and synchronization.

#### Nuclei Extraction and Library Preparation

After the final bleaching step, embryos were collected for nuclei isolation following the protocol described by Ooi *et al.* [81]. Nuclei quality was assessed by DAPI staining, and concentrations were determined by the mean of three independent counts using a Countess II FL automated cell counter. Concentrations were adjusted to meet the specific input requirements for the Active Motif ATAC-seq Kit (cat. no. 53150). Final libraries were sequenced on an Oxford Nanopore MinION flow cell using the Ligation Sequencing Protocol.

### AI usage

Large-language models Gemini 3.1 Pro (Google) and ChatGPT-5.3 Pro (OpenAI) were used to assist in writing the TBE model code; Gemini, ChatGPT, and Grammarly (v1.5.81) were used to proofread the manuscript for clarity.

## Acknowledgments

We thank A. Fire and A. Velazquez for discussions, O. Hobert and A. Fire for comments on the manuscript, and the researchers who made the public datasets used in this study accessible.

## Author contributions

F.A.: Conceptualization, Formal analysis, Investigation, Methodology, Visualization, Writing – original draft. M.P., H.A.: Investigation. S.E.M.: Investigation. S.H.: Supervision. C.F.-J.: Conceptualization, Supervision, Investigation, Writing – review and editing.

## Data availability

All data supporting the findings of this study are available within the paper and its Supplementary Information files. Annotated GenBank files for all transgenes generated in this study are provided as Supplementary Data 1. Long-read ATAC-seq data have been deposited in the NCBI Sequence Read Archive (SRA) under BioProject ID PRJNA1426581. Publicly available genomic datasets used in this study were obtained from WormBase (WS290), Dfam, Repbase, and previously published studies as cited in the Methods.

## Code availability

The Topological Bending Energy (TBE) model source code, along with the complete computational analysis pipeline used to process data and generate figures, is publicly available on GitHub at https://github.com/fffamk/TBE.

## Supplementary information

**Figure S1:**
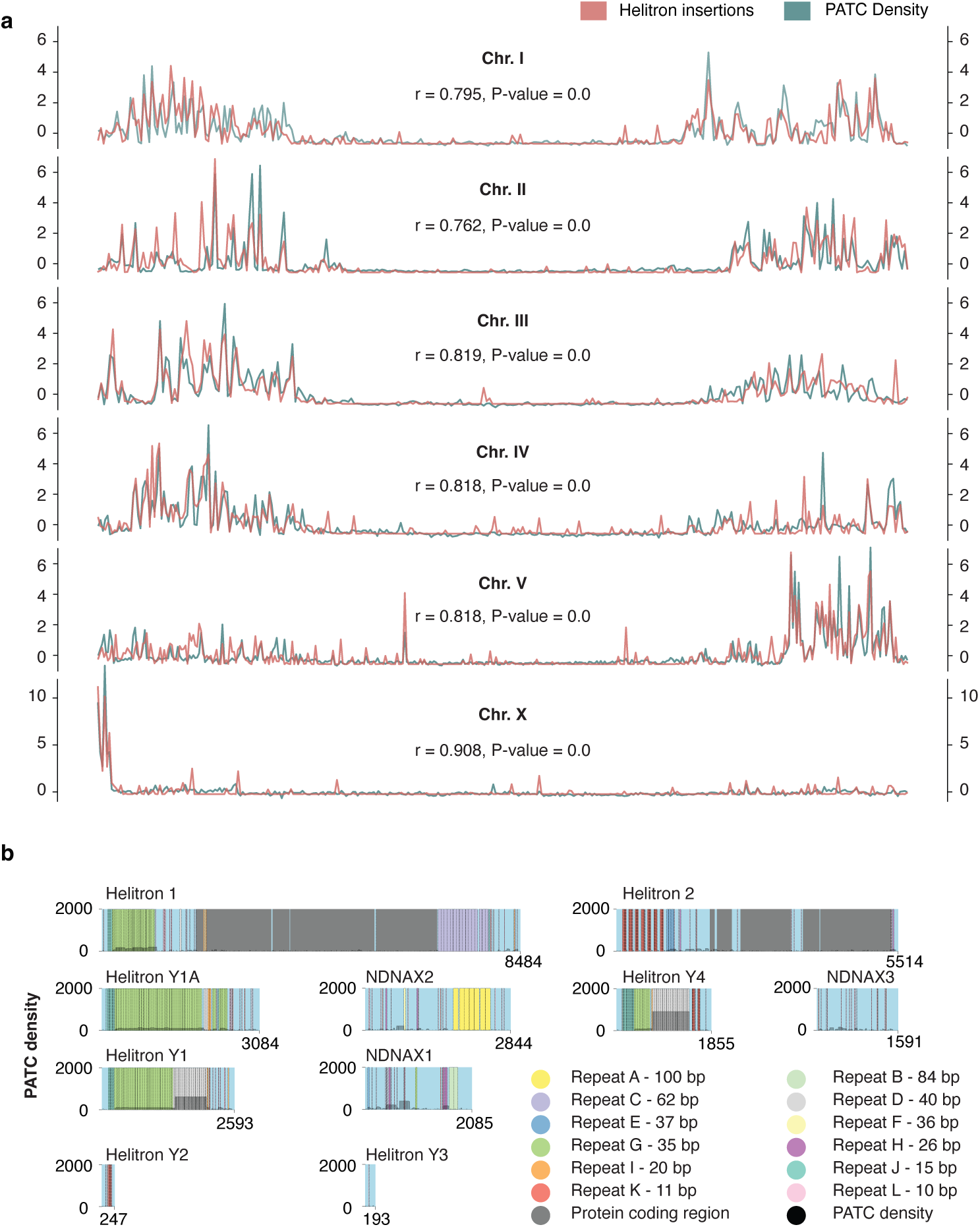
Genomic PATC signal correlates with Helitron insertions but is absent from ancestral consensus sequences. **a**, Genome-wide correlation of PATC density (Z-score, left axis) and Helitron insertion density (Z-score, right axis). **b**, PATC density across consensus sequences of the most abundant Helitron families in *C. elegans*. Colored blocks indicate internal minisatellite repeats.

**Figure S2:**
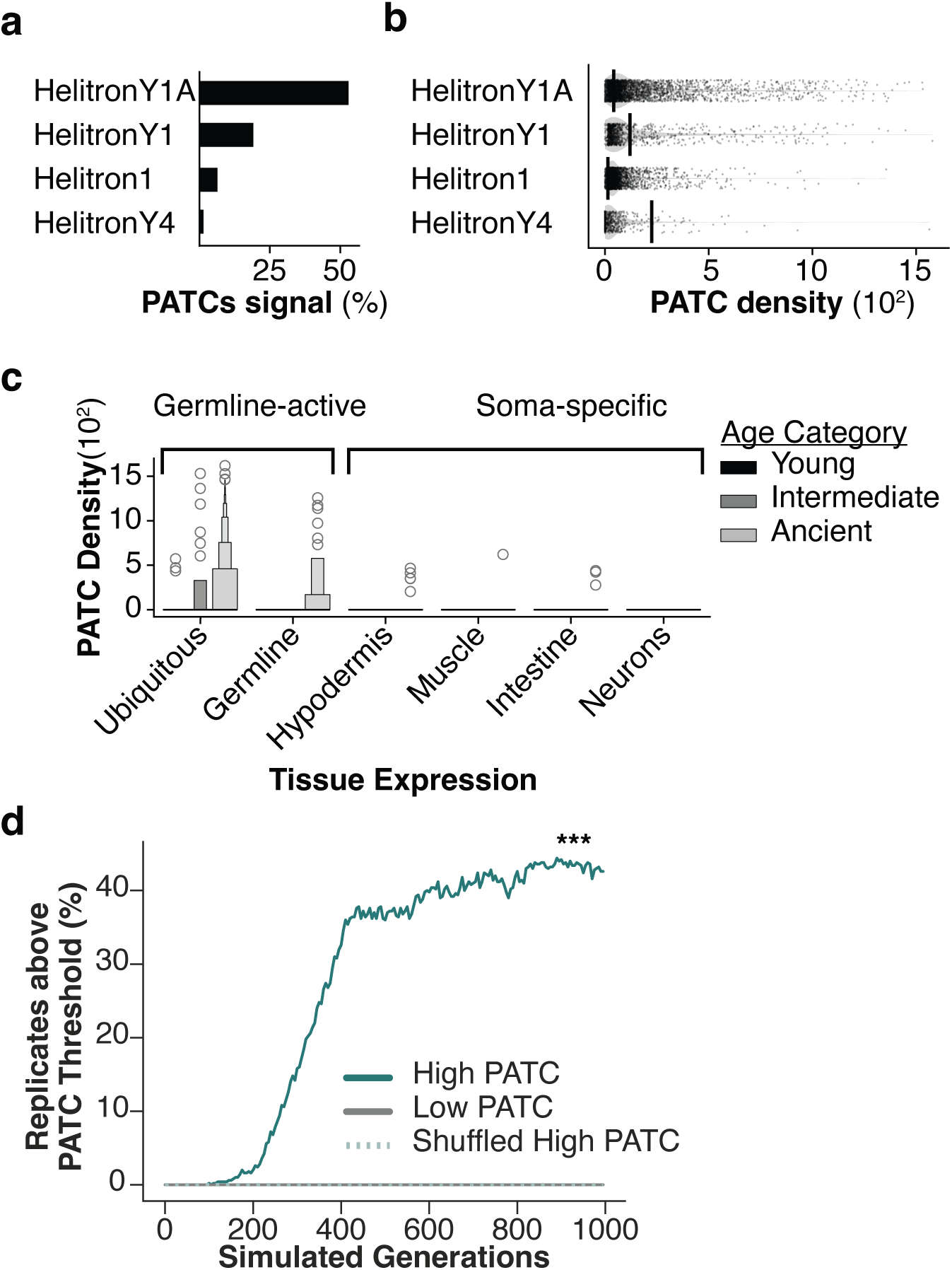
PATC evolution is specific to the germline. **a**, Contribution of specific Helitron families to the total PATC signal **b**, PATC density of individual genomic insertions (points) compared to their ancestral consensus sequences (vertical lines, right). **c**, PATC density of minisatellites stratified by evolutionary age and host gene tissue expression. **d**, Forward evolutionary simulation of PATC density over 1000 generations under high PATC (teal), low PATC (gray), or shuffled high PATC (dashed) (N = 500 replicates). Lines show percentage of replicates with PATC density > 95 (P < 0.001, pairwise Chi-squared test, final generation).

**Figure S3:**
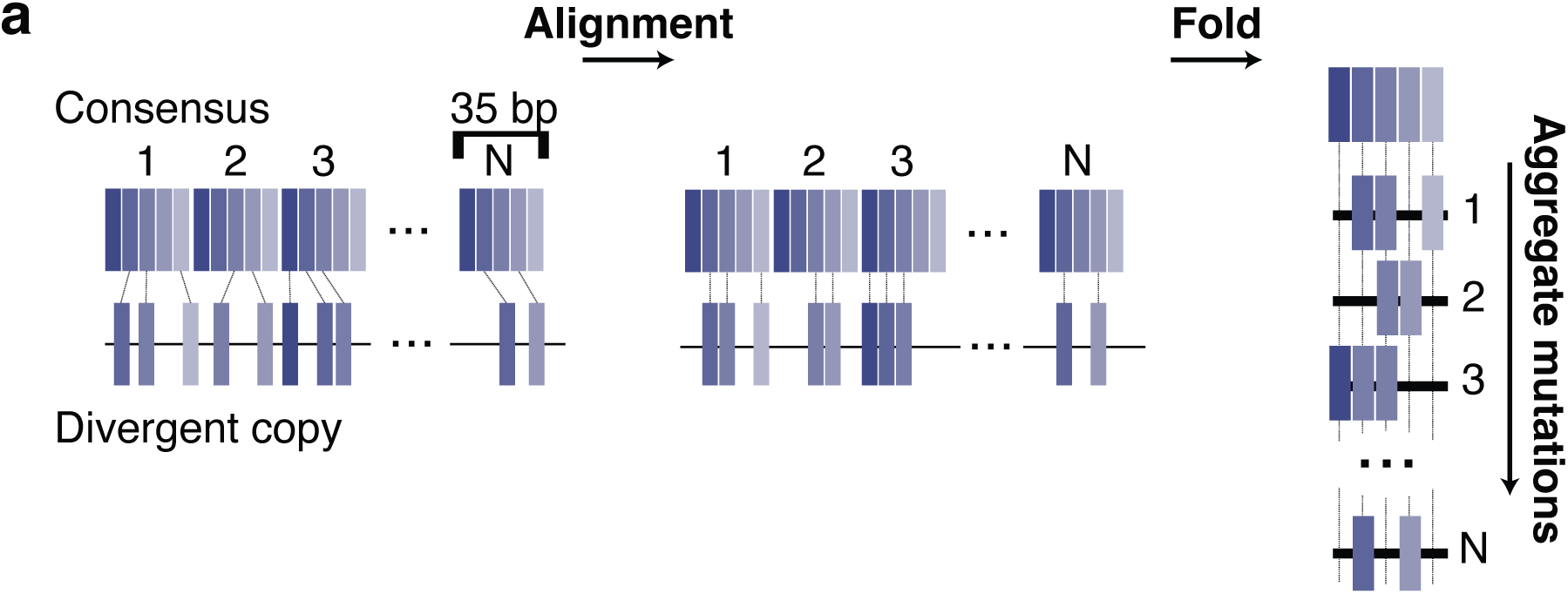
Monomer-folding alignment strategy. Schematic of the bioinformatics pipeline used to calculate position-specific mutability. Genomic minisatellite sequences are first aligned to the full-length Helitron consensus. The consensus is then decomposed into its constitutive 35-bp repeat units (monomers). Finally, all individual monomers—and their associated mutations— are mapped (“folded”) onto a single common 35-bp coordinate system, allowing substitutions from thousands of independent loci to be aggregated into a single mutability profile.

**Figure S4:**
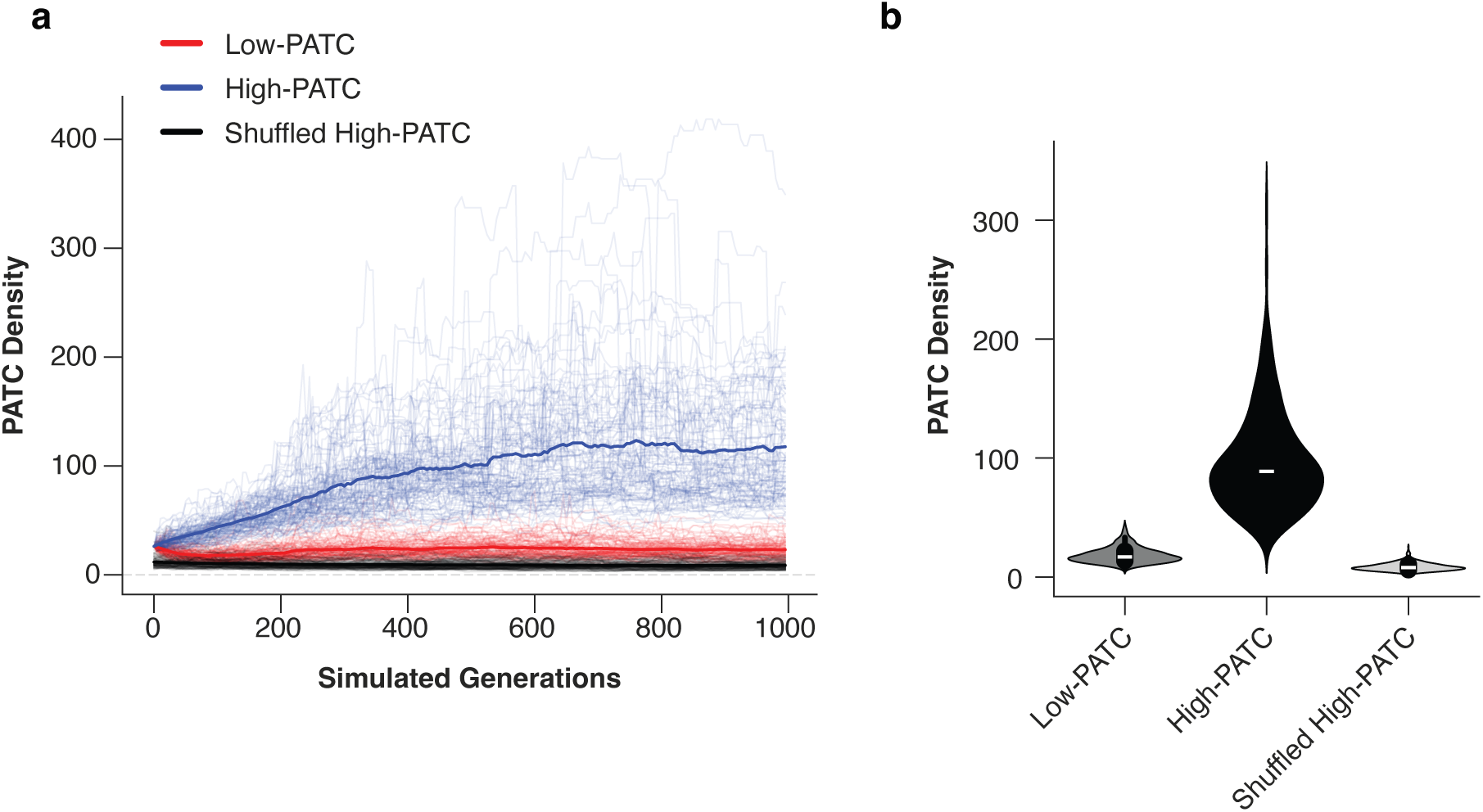
Mutational bias robustly drives PATC evolution. **a**, PATC density trajectories for individual replicates (N = 500) over 1000 simulated generations under high PATC, low PATC, and shuffled models. **b**, Distribution of PATC density at final generation (1000).

**Figure S5:**
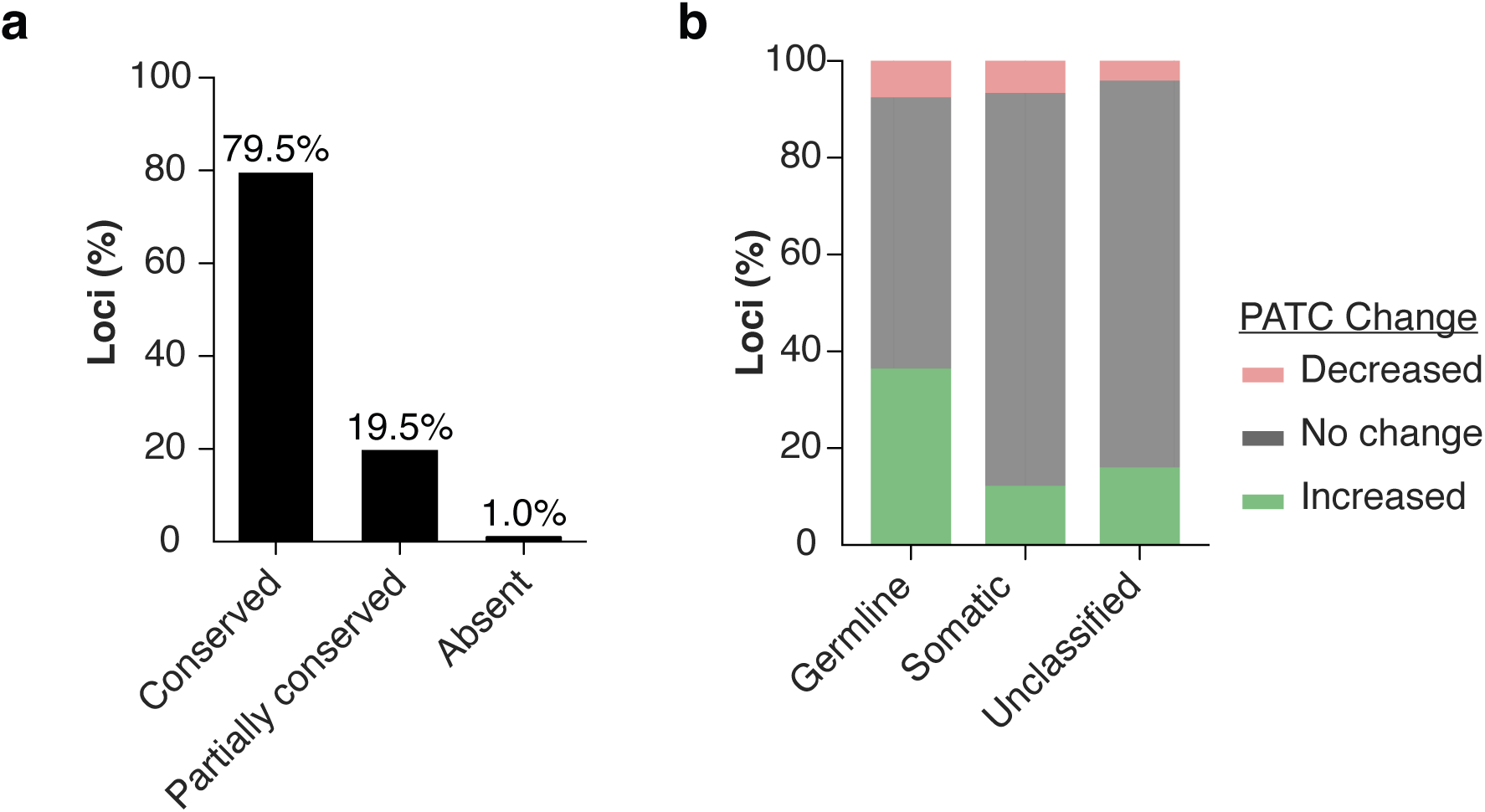
Ongoing germline-specific evolution of PATCs in wild populations. **a**, Conservation status of 44,305 Helitron-derived minisatellites across 17 wild *C. elegans* isolates. **b**, Change in PATC density for conserved loci relative to N2 reference, categorized by host gene expression (P < 0.005, Chi-squared test).

**Figure S6:**
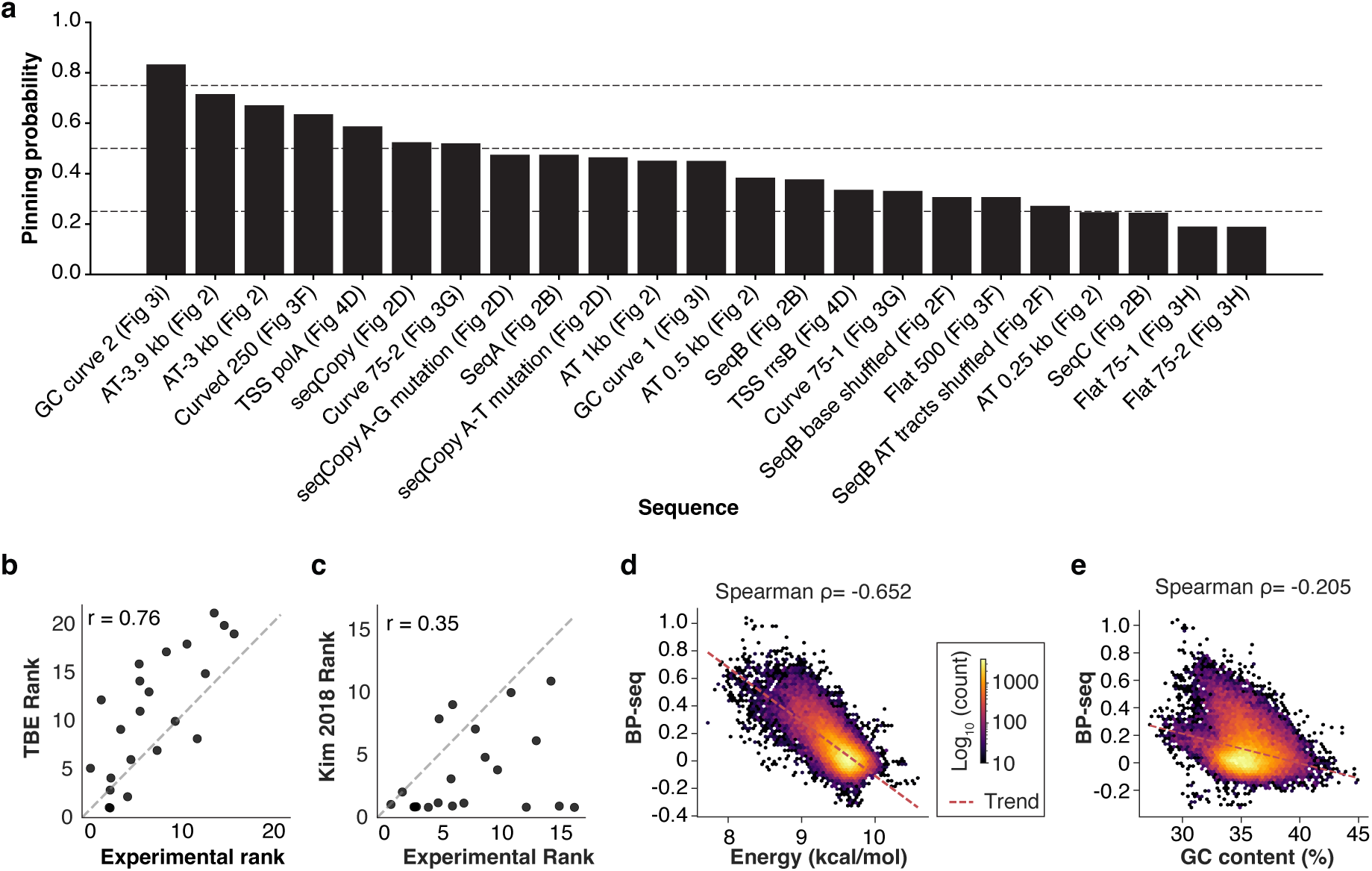
TBE model accurately predicts *in vitro* pinning and *in vivo* supercoiling. **a**, Predicted pinning probability calculated using the TBE model for each DNA construct assayed in Kim *et al*. [28] ordered by predicted probability. **b**, Rank–rank comparison between TBE model predictions and experimental pinning ranks (r = 0.76). **c**, Rank–rank comparison between the Kim *et al*. predictions and experimental ranks (r = 0.35). **d**, Scatterplot comparing BP-seq and TBE energy predictions (3kb bin mean; Spearman’s ρ = −0.65). **e**, Scatterplot comparing BP-seq and GC content (3kb bin mean; Spearman’s ρ = −0.21).

**Figure S7:**
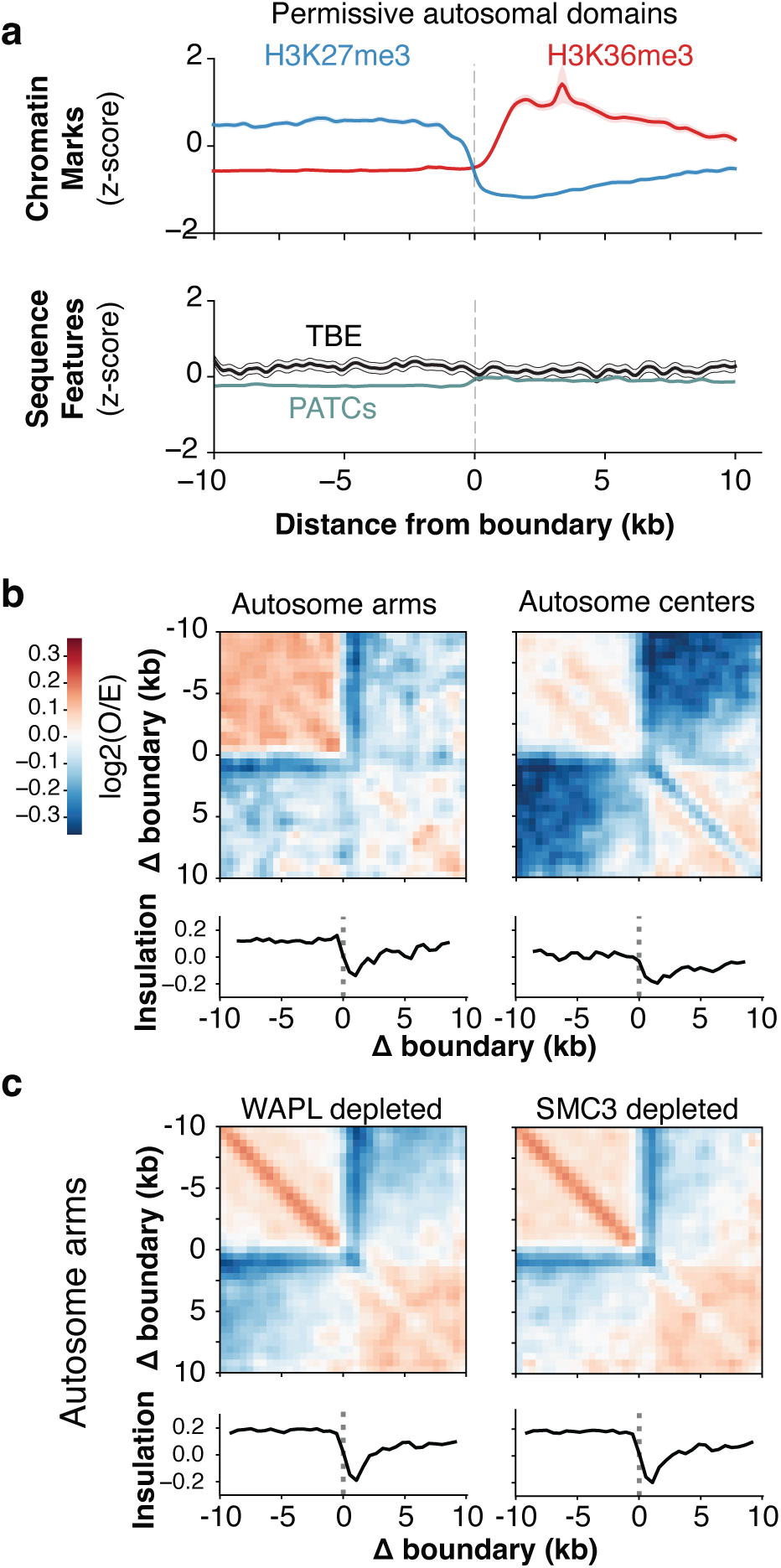
Autosomal centers are not enriched for PATCs or low TBE. **a**, Same analysis as Fig. 4a but centered on boundaries within the central part of the autosomes. (**b–c**) Previously published Hi-C [38, 39] log_2_(observed/expected) pileup view of the same boundaries in Fig. 4a and under somatic auxin-induced depletion of WAPL (left) and SMC3 (right).

**Figure S8:**
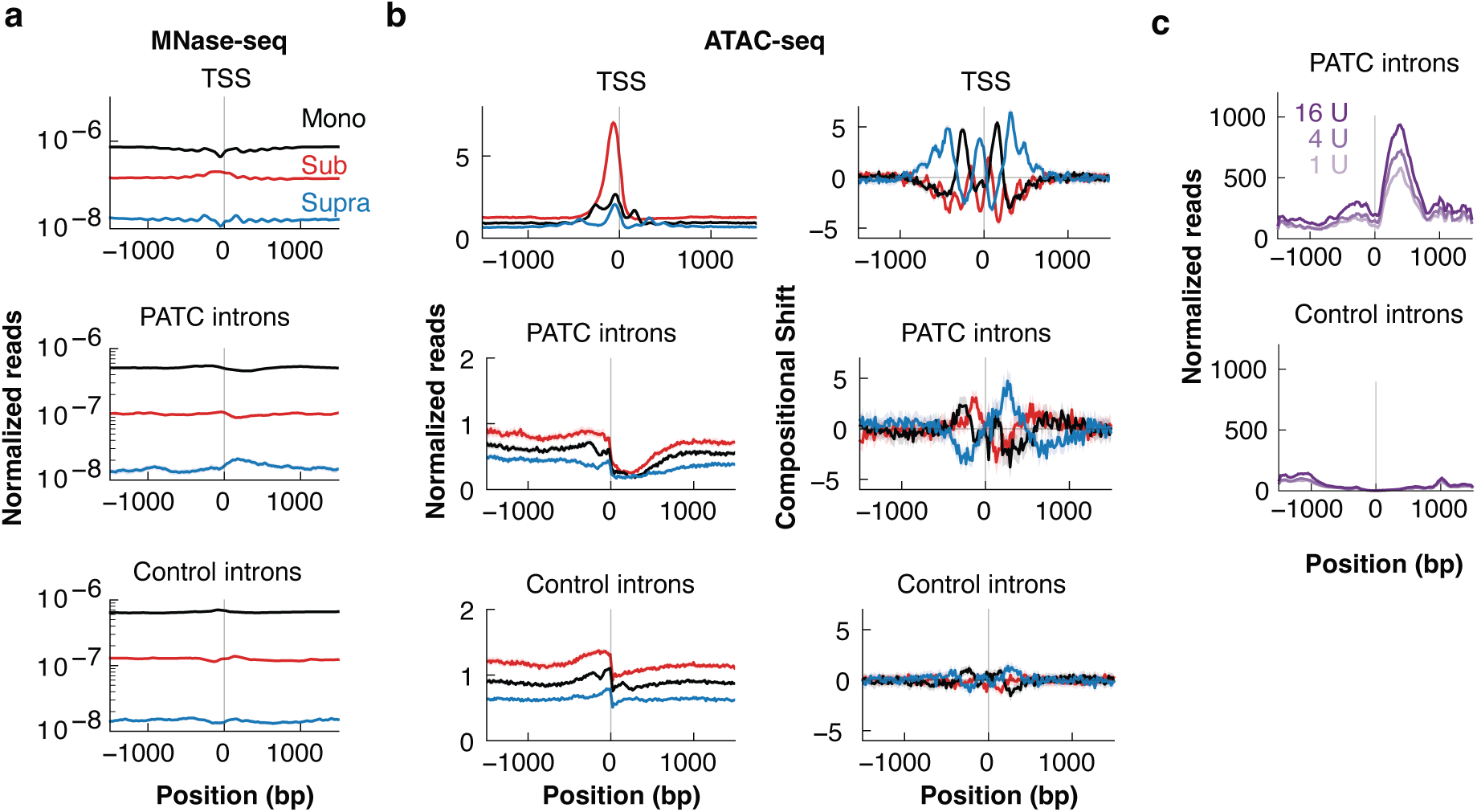
Fragment analysis validates PATC chromatin structure. **a**, Previously published MNase-seq [41] fragment coverage centered on TSS, PATC-rich introns, and matched control introns, similar to Fig. 5a. **b**, Previously published ATAC-seq [41] fragment coverage and compositional shift stratified by size: sub-nucleosomal (< 135 bp), mono-nucleosomal (150– 260 bp), and supra-nucleosomal (> 300 bp). **c**, Supra-nucleosomal fragment enrichment versus previously published MNase concentrations [41] for PATC-rich (top) and control (bottom) introns.

**Figure S9:**
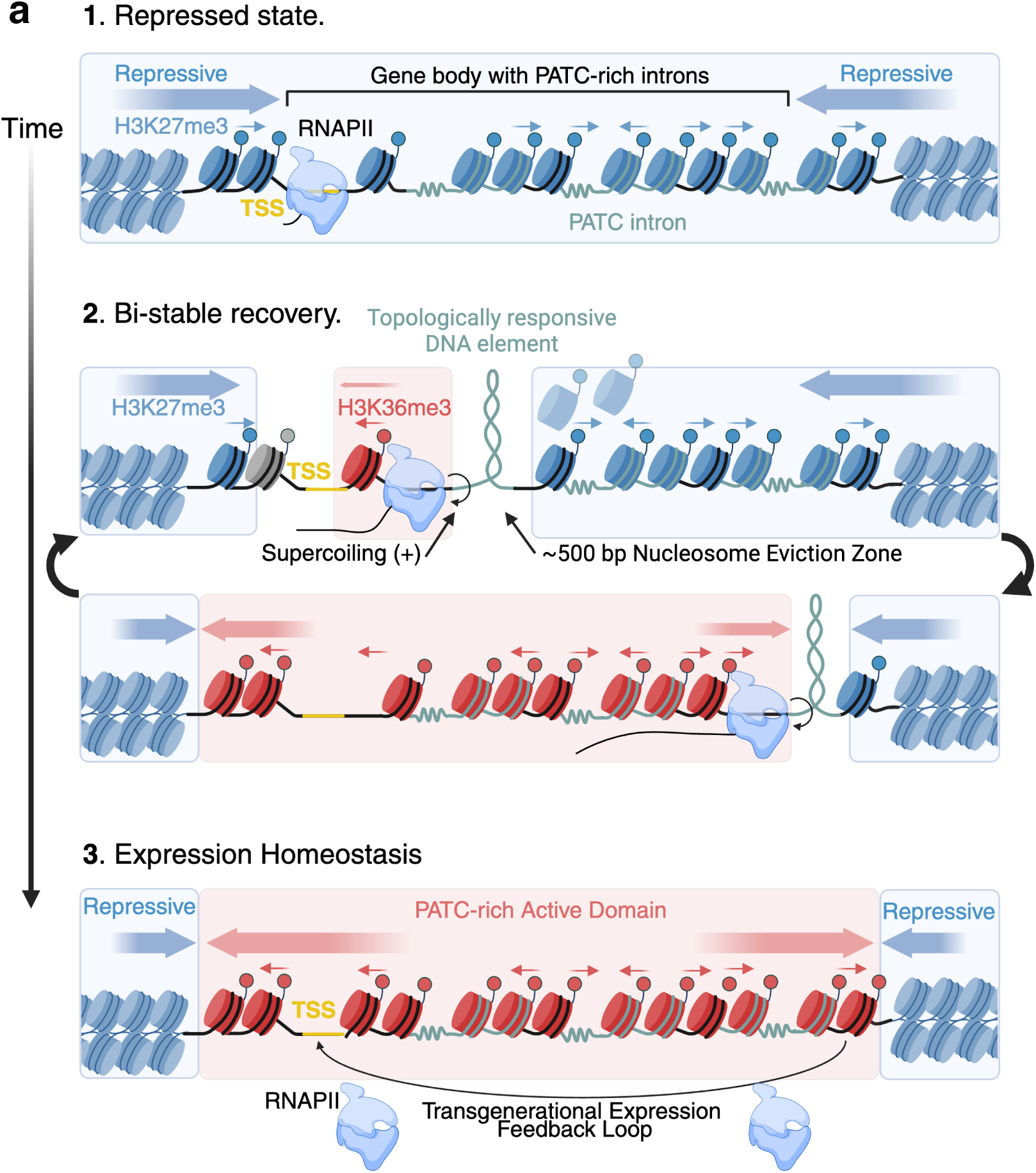
Model of mechanical chromatin insulation and the transgenerational establishment of gene expression homeostasis. **a**, **1. Repressed State:** A silenced locus is coated by repressive H3K27me3 nucleosomes. The intrinsic curvature of PATC-rich introns encodes baseline topological stress (zig-zag lines). **2. Bi-stable Recovery:** Following silencing trigger removal, initial RNAPII transcription introduces positive torsional stress. PATCs pin this torsion into a topologically responsive DNA element (TRDE), physically evicting repressive nucleosomes within a ∼500 bp zone. Incomplete transcriptional read-through creates a transgenerational bi-stable state, but PATC barriers enable significantly faster recovery than controls. **3. Expression Homeostasis:** Successive RNAPII elongation drives active H3K36me3 deposition, fully clearing repressive marks. PATC-pinned TRDEs act as persistent physical boundaries against flanking H3K27me3 domains, securing a stable, permissive chromatin state. Created with https://BioRender.com.

**Figure S10:**
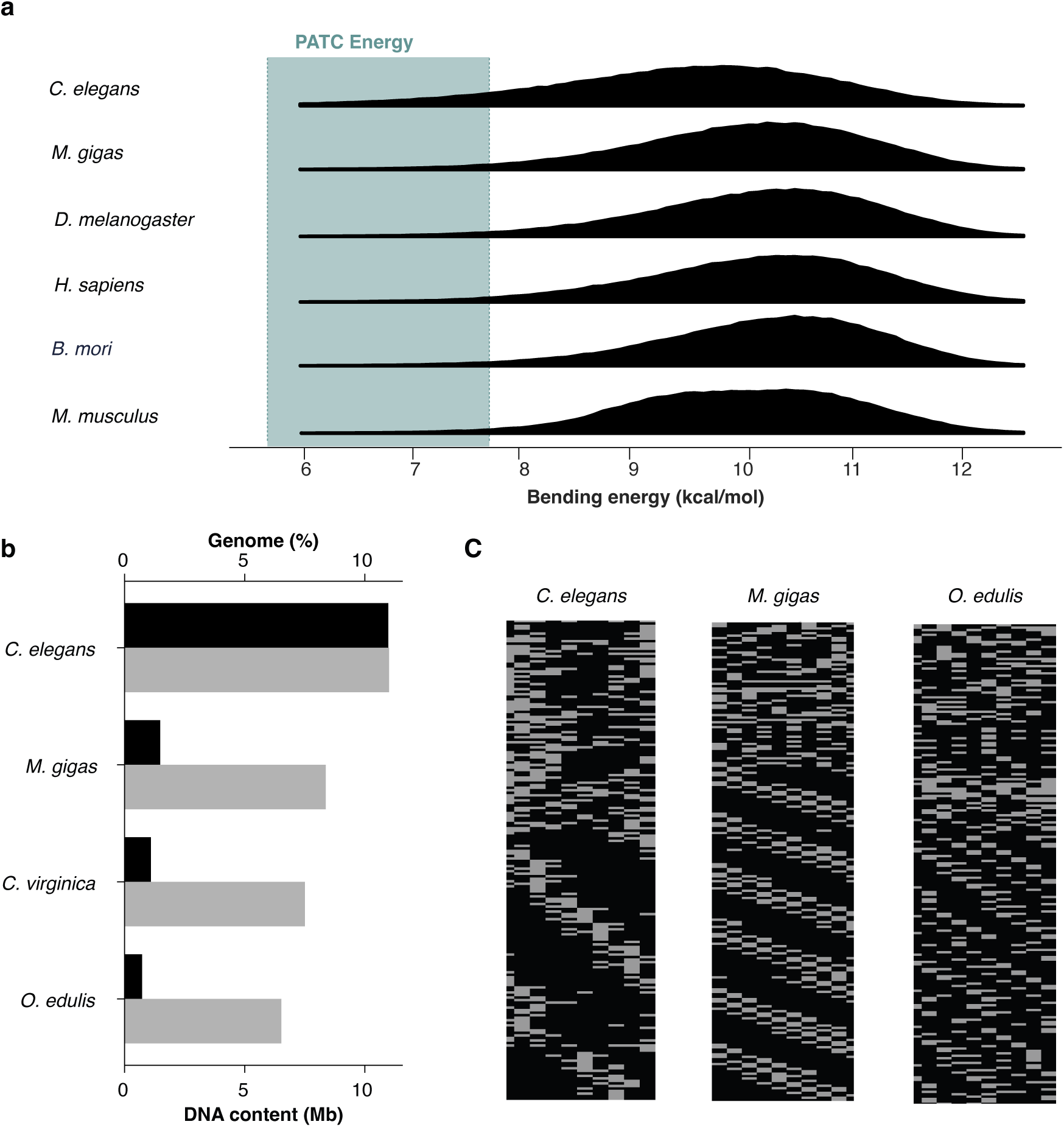
Convergent evolution of PATC-like elements across Metazoa. **a**, Distribution of genomic bending energies. Histograms representing the TBE landscape for representative metazoan genomes. The shaded teal region (“PATC Energy”) denotes the energetic threshold characteristic of *C. elegans* PATCs, highlighting the presence of high-pinning sequences across divergent phyla **b**, Genomic distribution of PATCs. Comparative analysis of high-density periodic sequences in *C. elegans* and three representative oyster species: the Pacific oyster (*M. gigas*), the Eastern oyster (*C. virginica*), and the European flat oyster (*O. edulis*). Sequences were identified using the Fire et al. algorithm (threshold: 60 PATC density). Grey bars represent the absolute cumulative size of PATC-like sequences (Mb); black bars denote their relative fraction of the total genome (%). **c**, Nucleotide-level periodicity. Visualization of primary sequence architecture for representative PATC-like elements in *C. elegans*, *M. gigas*, and *O. edulis*. Each column represents a 10-bp interval; grey boxes indicate A/T bases and black boxes indicate G/C bases, illustrating the 10-bp periodicity of A-tracts characteristic of pinning sequences across divergent species.

**Figure S11:**
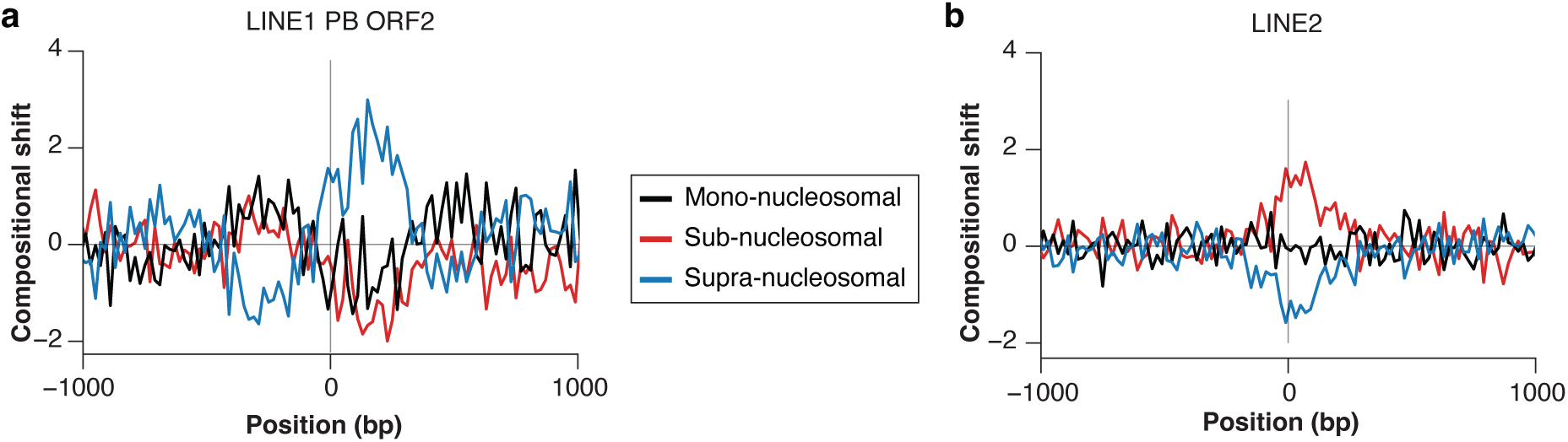
ATAC-seq fragment analysis of LINE-1 and LINE-2. (**a–b**) Meta-profiles of previously published ATAC-seq [53]. Fragment coverage centered on human LINE-1 (**a**) and the extinct LINE-2 (**b**) families. Reads are stratified by size into sub-nucleosomal (< 120 bp, red), mono-nucleosomal (170–230 bp, black), and supra-nucleosomal (> 300 bp, blue) classes. Note the presence of the diagnostic “supra-nucleosomal” footprint in active LINE-1 elements (**a**), indicating a compact, non-nucleosomal topological structure that is absent in the LINE-2 lineage (**b**).

